# PFAS assessment in fish – samples from Illinois waters

**DOI:** 10.1101/2023.08.29.555412

**Authors:** Mia Sands, Xing Zhang, Tor Jensen, Michael La Frano, Mindy Lin, Joseph Irudayaraj

## Abstract

Per- and Polyfluoroalkyl substances (PFAS) have been widely used in various industries, including pesticide production, electroplating, packaging, paper making, and the manufacturing of water-resistant clothes. This study investigates the levels of PFAS in fish tissues collected from four target waterways (15 sampling points) in the northwestern part of Illinois during 2021-2022. To assess accumulation, concentrations of 17 PFAS compounds were evaluated in nine fish species to potentially inform on exposure risks to local sport fishing population via fish consumption. At least four PFAS (PFHxA, PFHxS, PFOS, and PFBS) were detected at each sampling site. The highest concentrations of PFAS were consistently found in samples from the Rock River, particularly in areas near urban and industrial activities. PFHxA emerged as the most accumulated PFAS in the year 2022, while PFBS and PFOS dominated in 2021. Channel Catfish exhibited the highest PFAS content across different fish species, indicating its bioaccumulation potential across the food chain. Elevated levels of PFOS were observed in nearly all fish, indicating the need for careful consideration of fish consumption. Additional bioaccumulation data in the future years is needed to shed light on the sources and PFAS accumulation potential in aquatic wildlife in relation to exposures for potential health risk assessment.

## 1. Introduction

Per- and Polyfluoroalkyl substances (PFAS) are a class of organic compounds wherein the hydrogen atoms in the carbon chain of aliphatic hydrocarbons are replaced by fluorine atoms (Evich et al., 2022). Due to their high C-F bond energy, chemical stability, and hydrophobicity, PFAS have found widespread use in industries such as pesticide production, electroplating, packaging, paper manufacturing, water-resistant clothing manufacturing, and electronics. Due to its prolonged environmental half-life, bioaccumulation potential, and capacity for long-range transport, PFAS are classified as persistent organic pollutants (Lallas, 2001; UNEP, 2019; Zhang et al., 2022). Exposure to PFAS compounds have raised significant concerns due to its potential impact on human and animal health (Solan et al., 2023; Wen et al., 2022). The U.S. Environmental Protection Agency (EPA) has established guidelines and standards for select PFAS in drinking water and/or soil in some U.S. states and internationally (EPA, 2016; ITRC, 2024). With the goal of phasing out the use of PFAS, Perfluorooctane sulfonate (PFOS) and Perfluorooctanoic acid (PFOA) were included in the Stockholm Convention (SC) as Persistent Organic Pollutants in 2009 and 2019, respectively (UNEP, 2019). Industries in the United States and in some parts of the world are voluntarily restricting the use of these chemicals banned by the SC (Gaines, 2023). As PFOS and PFOA use and production has decreased, production of PFAS chemicals with shorter carbon backbones has increased (Loganathan and Wilson, 2022).

PFAS are broadly categorized into two groups: long-chain and short-chain PFAS. Long-chain PFAS, exemplified by PFOS and PFOA, typically contain at least 8 carbon atoms. In contrast, shorter-chain PFAS, including Perfluorohexane Sulfonic Acid (PFHxS), Perfluorobutane sulfonic acid (PFBS), and Perfluorobutyrate Acid (PFBA), consist of less than 8 carbon atoms (Chambers et al., 2021). Further classification distinguishes ultra short-chain and short-chain PFAS, with 2-3 and 4-7 fully fluorinated carbon atoms, respectively (Ateia et al., 2019; Wang et al., 2017). Generally, long-chain PFAS tend to partition to sediments, whereas shorter chain compounds typically remain predominantly in the dissolved state. It is important to note that while longer chain compounds often accumulate in sediments, which inturn can also act as a potential source of these compounds to both surface and groundwater (Labadie and Chevreuil, 2011).

The remarkable chemical stability of PFAS contributes to its prolonged persistence in the environment, allowing for long-distance migration through various mechanisms, including water flow and atmospheric conversion, reaching even remote locations such as the North and South Pole (Yeung et al., 2017). Extensive research has investigated the multi-agency distribution and ecological impacts of long-chain PFAS such as PFOS and PFOA (Anderko and Pennea, 2020; Rahman et al., 2014). Pham et al. reported that PFOS and PFOA in the surface water of the Dianchuan Basin in Japan, reached up to 123 ng/L and 260 ng/L, respectively (Lein et al., 2008). U.S. EPA noted that PFOS and PFOA in soils are likely to be a major source of PFAS contamination (Renner, 2009). In Australia, drinking water samples have shown PFOS and PFOA levels at 16 ng/L and 9.7 ng/L, respectively (Thompson et al., 2011). PFAS testing in 19 major water systems in China revealed both short-chain Perfluorobutanesulfonic acid (PFBS) and PFOA (Wang et al., 2016). In the Great Lakes of North America, the cyclic perfluorinated acid, perfluoroethylcyclohexanesulfonate, was detected at concentrations similar to PFOS (De Silva et al., 2011). While long-chain PFAS has been the primary focus of active research, the production and application of short-chain PFAS continue to rise, leading to their widespread accumulation in aquatic systems. Despite the discontinuation of the use of certain PFAS within the U.S., imported sources from other parts of the world persist and concerns exist on the use of short-chain replacements (De Silva et al., 2021; Pickard et al., 2022). Short-chain PFAS are known to be less absorbent, more persistent, and more mobile than long-chain PFAS (Li et al., 2020; Mumtaz et al., 2019). The potential health risks associated with short-chain PFAS are thought to be as high as the long-chain PFAS (Brendel et al., 2018; Houtz et al., 2013; Li et al., 2020).

A significant knowledge gap remains in understanding the health impact of these emerging contaminants in both marine mammals and humans through fish consumption. PFAS compounds tend to accumulate in humans, leading to various adverse effects. PFOS and long-chain perfluoroalkyl carboxylates (PFCAs), such as perfluoroundecanoic acid (PFUnA; C11), can accumulate in tissues (Labadie and Chevreuil, 2011). Furthermore, PFOS and PFOA were detected in serum samples from 105 Danish men, with levels negatively correlated with sperm count and quality (Joensen et al., 2009). PFOA has been linked to reproductive toxicity, immunotoxicity, and has been classified as a possible carcinogen by the World Health Organization’s International Agency for Research on Cancer (Cordner et al., 2019; Humans, 2017; Shane et al., 2020; Wen et al., 2020a; Wen et al., 2020b). PFOS has also been associated with multiple toxic effects, including developmental and reproductive issues, as well as hepatotoxicity (Rashid et al., 2023; Wen et al., 2022).

While the general population is exposed to low levels of PFAS from various sources such as food packaging and consumer products, individuals can be exposed to higher levels due to the consumption of contaminated food or drinking water (Fujii et al., 2015; Post et al., 2012). Numerous studies have also reported the presence of PFAS in the environment and wildlife (Barbo et al., 2023; Evich et al., 2022; Fair et al., 2019). In general, food and water seems to be the primary non-occupational exposure route for PFAS, with seafood containing the highest concentrations of PFAS (Augustsson et al., 2021; Denys et al., 2014; Domingo and Nadal, 2017). Fish consumption has experienced a notable increase, rising by roughly 30% in the U.S. over recent decades (Loke et al., 2012). In measurements of the top predator lake trout (*Salvelinus namaycush*) in the Great Lakes, PFOS was the predominant PFAS observed, with concentrations as high as 121 ± 14 ng/g (Furdui et al., 2007), along with PFOA levels ranging from 0.65–5.5 ng/L (De Silva et al., 2011). Among the PFAS detected, PFOS is generally the most prevalent and found in the highest concentrations in fish, although other PFAS are also frequently detected (Houde et al., 2011). Studies such as those by Delinsky et al. (Delinsky et al., 2010) in Minnesota lakes and several Upper Mississippi River locations, and Fair et al. (Fair et al., 2019) in the Charleston Harbor and tributaries in South Carolina, have reported PFAS in various aquatic ecosystems. The reported levels in the Great Lakes (Stahl et al., 2023) were generally lower than those in the Minnesota River (Lin et al., 2021), and substantially lower than those in urban rivers in America (Coy et al., 2022; Goodrow et al., 2020a; MacGillivray, 2021). However, there is limited to no information on PFAS contamination in fish from waterways across Illinois. To conduct a comprehensive risk analysis, it is necessary to obtain data on PFAS levels by its type and species from specific rivers and tributaries.

The objectives of this study were to: 1) conduct an initial assessment of the occurrence of 17 PFAS, both long- and short-chain, and their distribution across nine fish species from local Illinois rivers collected in 2021 and 2022, 2) determine the potential risks of PFAS exposure in these species, and 3) estimate the dietary PFAS exposure due to fish consumption. The findings of this study could shed light on the types of PFAS compounds in fish obtained from waterways around the state. In the long-term with periodic sampling, this information could aid local agencies in efficient management/mitigation of human exposure risks.

## 2. Materials and Methods

### 2.1 Chemicals and reagents

MPFAC-2ES, a mixture containing 19 mass-labelled standards containing C_4_-C_14_ PFCAs, C_4_-C_10_ PFSAs, FOSA, N-MeFOSAA, N-EtFOSAA, as well as 4:2, 6:2, and 8:2FTS was purchased from Wellington Laboratories (Canada). Target compounds included four perfluorocarboxylic acids (PFBA, PFHxA, PFOA and PFDA) and three perfluorosulfonic acids (PFBS, PFHxS and PFOS). These compounds were purchased from Wellington Laboratories (Canada) and SynQuest Laboratories (USA). Detailed information is provided in Table S1. The stock solutions were prepared in methanol and kept in polypropylene (PP) tubes at 4°C. HPLC-grade methanol, with a purity of at least 99.9%, from Sigma-Aldrich (USA), was used as a solvent for HPLC analysis.

### 2.2 Study area

Based on their proximity to potential sources of PFAS and the likelihood of recreational and subsistence fishing, samples were collected from Rock River, Pecatonica River, Sugar River, and Yellow Creek in Illinois (see **Figure 1** and Table S2). Rock River is the largest of the waterways sampled flowing through larger urban centers with the Pecatonica River as a smaller tributary. Sugar River and Yellow Creek are small tributaries to the Pecatonica River. The four waterways/aquatic systems are in the northwest corner of Illinois.

**Figure 1.**
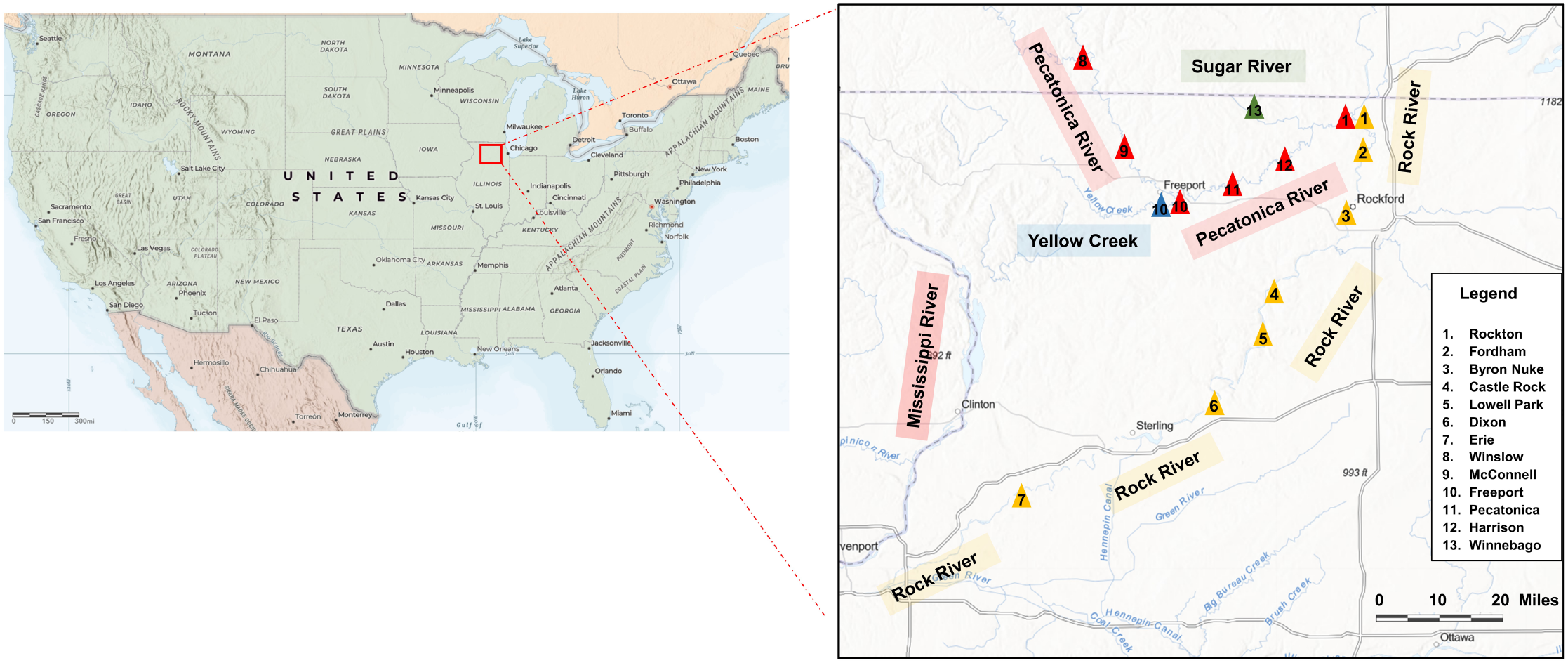
Map of the 15 sampling sites located in four major rivers in Illinois, USA during the years 2021 and 2022. Noted that fish samples were only collected from sampling sites 1-7 along the Rock River in 2021.

The Rock River is a significant waterway in the state of Illinois. It stretches for approximately 299 miles, flowing through various cities and towns. The Rock River is also one of Mississippi River’s major tributaries and it merges near the Quad Cities region of Illinois and Iowa. It has served as a vital transportation route for industries, commerce, and recreational activities and facilitated trade and transportation of goods, supports various species of fish, birds, and other wildlife. The watersheds have varying land-use and land-cover compositions, ranging from agricultural dominance and forest dominance to heavy urbanization. Sampling activities were conducted during the period between October to November 2021 and July to November 2022 to collect relevant data and information.

### 2.3 Sample collection

A total of 52 fish samples were collected from the Rock River in 2021 for fillet contaminant analysis. Five fish species were collected in 2021: Bluegill (*Lepomis macrochirus*), Common Carp (*Cyprinus carpio*), Smallmouth Bass (*Micropterus dolomieu*), Walleye (*Sander vitreus*), and Channel Catfish (*Ictalurus punctatus*). Nine fish species with a total of 70 fish samples were collected in 2022 from four waterways. The additional fish species collected in 2022 were Sauger (*Sander canadensis*), Smallmouth Buffalo (*Ictiobus bubalus*), Black Crappie (*Pomoxis nigromaculatus*) and Northern Pike (*Esox Lucius*). These species were selected to represent both pelagic and benthic environments and encompass various trophic levels. The demographic information can be found in **Table 1**.

**Table 1.** Physical characteristics and demographics across all samples

| Number | Fish Code | Species | Latin Name | Habitat | Trophic level descriptors | Trophic level | Length(mm) | Weight(g) |
| --- | --- | --- | --- | --- | --- | --- | --- | --- |
| 1 | BLG | Bluegill | Lepomis macrochirus | Benthic | Primary consumer (herbivore) | 2 | 173 ± 28 | 126 ± 51 |
| 2 | SAB | Small-mouth Buffalo | Ictiobus bubalus | Benthic | Primary consumer (herbivore) | 2 | 420 | 1080 |
| 3 | CAP | Common Carp | Cyprinus carpio | Benthic | Primary consumer (omnivore) | 2 | 579 ± 99 | 2933 ± 1569 |
| 4 | SAR | Sauger | Sander canadensis | Benthic | Lower Trophic Level Insectivore/Piscivore | 3 | 439 ± 58 | 797 ± 449 |
| 5 | SMB | Small-mouth Bass | Microp-terus dolomieu | Benthic | Lower Trophic Level Insectivore/Piscivore | 3 | 321 ± 66 | 502 ± 288 |
| 6 | WAE | Walleye | Sander vitreus | Benthic | Lower Trophic Level Insectivore/Piscivore | 3 | 419 ± 108 | 850 ± 555 |
| 7 | BLC | Black Crappie | Pomoxis nigromaculatus | Pelagic and Benthic | Lower Trophic Level Insectivore/Piscivore | 3 | 235 ± 24 | 216 ± 75 |
| 8 | CCF | Channel Catfish | Ictalurus punctatus | Benthic | Top Trophic Level Insectivore/Piscivore | 4 | 483 ± 73 | 1111 ± 598 |
| 9 | NOP | Northern Pike | Esox lucius | Benthic | Top Trophic Level Insectivore/Piscivore | 4 | 710 | 2145 |

Fish samples were collected by the Illinois Department of Natural Resources (IL-DNR). To ensure sample integrity, each fish was individually wrapped in thick aluminum foil and then placed inside a sealed polyethylene bag. The samples were kept on ice for 3-6 hours before being frozen at -20°C. They remained frozen until they were ready for preparation and subsequent analysis.

### 2.4 Sample extraction and Instrumental analysis

Sample processing steps were adapted from other works (Fair et al., 2019; Valsecchi et al., 2021). In brief, methanol was used to extract PFAS from the fish fillet sample. Once the Mass-labelled PFAS extraction standard solution containing 19 components (MPFAC-2ES) was added, PFAS were measured through liquid chromatography-mass spectrometry (LC-MS) using both internal and external standards (Table S8 and S9), with calibration done at various concentration levels. LC-MS was provided by the Carver Metabolomics Core Facility of the Roy J. Carver Biotechnology Center, University of Illinois Urbana-Champaign (Genualdi and deJager; Shoemaker and Tettenhorst). Using an Agilent 1260 Infinity II HPLC system (Agilent Technologies, Santa Clara, CA, USA), PFAS metabolites were separated with a Phenomenex Kinetex PS C18 100A (2.6μm, 100 x 4.6mm) column (Phenomenex, Torrance, CA, USA) via a gradient method consisting of mobile phase A: H2O+20mM ammonium acetate and mobile phase B: methanol running at a flow rate of 0.35 mL/min. The gradient was 0-2 min = 90% A; 2-10min = 0% A; 18.1-24min = 90% A. A Sciex 6500+ triple quadrupole MS (Sciex, Framingham, MA, USA) operating in negative ionization mode using multiple reaction monitoring (MRM) was used to screen for over 20 PFAS compounds.

### 2.5 Quality assurance and quality control (QA/QC)

The selection of materials used throughout sample processing and analysis was done with the intent of minimizing PFAS contamination. Materials such as polypropylene, glass, and metal were used. Metabolites were quantified with Sciex Analyst software using 5- to 7-point calibration curves adjusted for stable-isotope- and deuterium-labeled internal standards (Wellington Laboratories, Guelph, ON). A delay column (Halo PFAS delay column, 2.7 μm, 4.6 mm x 50 mm; Advanced Materials Technology, Wilmington, DE, USA) was placed between the mobile phase mixer and sample injector to momentarily trap any system related PFAS interferences due to tubing or mobile phase. Solvent and process blanks were periodically analyzed throughout the sequence to test for the presence of PFAS contamination. The solvent blanks included the same reconstitution solvent used for samples (70% methanol). The process blanks followed the entire process of extraction and analysis. QCs were run to assess reproducibility of quantitation at varying concentrations. Reference materials used included analysis of mixtures of PFAS standards. The analytical lab was blinded to the identity of the sample groups. All samples were randomized and analyzed in a single batch.

### 2.6 Risk assessment

In this study, the Risk Quotient (RQ) method was used to evaluate the ecological risk degree of PFAS in fish. RQ is the ratio of the measured environmental concentration (MEC, in ng/L) and the predicted non-effect concentration (PNEC, in ng/L) of the actual sample and calculated using the formula:

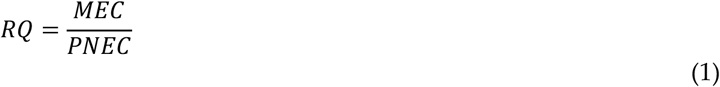

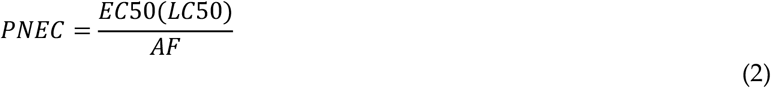

Where: PNEC was obtained from toxicological data. LC50 was the lowest 50% lethal concentration (ng/L) obtained by toxicological tests across organisms. EC50 was the minimum 50% effective concentration (ng/L) obtained by toxicological tests across organisms. AF is the EC50 value of the assessment factor (three trophic levels (algae, crustacea, fish)). Ecological toxicology data was obtained from the US EPA’s Integrated Risk Information System (IRIS) database at https://www.epa.gov/iris.

The risk stratification criteria for each PFAS are as follows:

RQ < 0.01, no risk;

0.01≤RQ < 0.1, low risk;

0.10≤RQ < 1.00, medium risk;

RQ≥1.00, high risk.

### 2.7 Fish consumption

Based on national and local fish consumption, we analyzed the impact of consuming fish on human exposure to PFAS. The daily trigger concentration (DTC) (Eq. (3)) is the ratio of Reference Dose (RfD; the daily dose in ng/kg/day expected without risk from lifetime exposure) multiplied by average body weight (BW, assuming an average body of 70 kg), then divided by meal size(g).

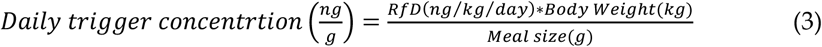

Where: Body weight = 70 kg; Meal size = 227 g.

## 3. Results and discussion

### 3.1 Concentration and composition of PFAS in fish by location

Data was collected from 122 fish samples spanning 2021 to 2022. Detailed findings on PFAS detected are outlined in Tables S6 and S7. In 2022, seventeen PFAS were assessed across Rock River, Pecatonica River, Sugar River, and Yellow Creek, while seven major PFAS were evaluated in fish from Rock River in 2021. Mean concentrations of six highly accumulated PFAS for both years are depicted in **Figure 2**. The mean percentage composition of different PFAS at sampling sites is illustrated in Figures S2a and S2e. **Figure 2a** shows that the highest concentration in 2021 is PFBS driven by samples collected downstream of Byron, home to a nuclear power plant (Table S6). Concentrations of PFBS in 2021 are followed by PFOS and PFHxS, compounds identified in surface water by multiple other studies (Ali et al., 2021; Barbo et al., 2023; Fair et al., 2019; Pickard et al., 2022). **Figure** S2d and S2e show that the highest concentration in 2022 is PFHxA, followed by PFOS among the seventeen PFAS. Among the four watersheds evaluated in 2022, sampling sites in Rock River exhibited the highest PFAS concentration, while smaller Pecatonica River tributary showed the lowest concentration and detected the fewest types of PFAS compounds **(Figure 2b)**. This disparity could be explained by the lesser industrial activities and urban runoff in the upstream areas of Pecatonica River. Correlations between each PFAS and contribution of each PFAS are shown in Figures S2b-c, S2f-g, Table S3-4. While frequent reports identify PFOA occurrence in fish tissue (Fair et al., 2019; Goodrow et al., 2020b; Langberg et al., 2022), this study only detected the compound in Rock River with none detected in samples from Pecatonica River.

**Figure 2.**
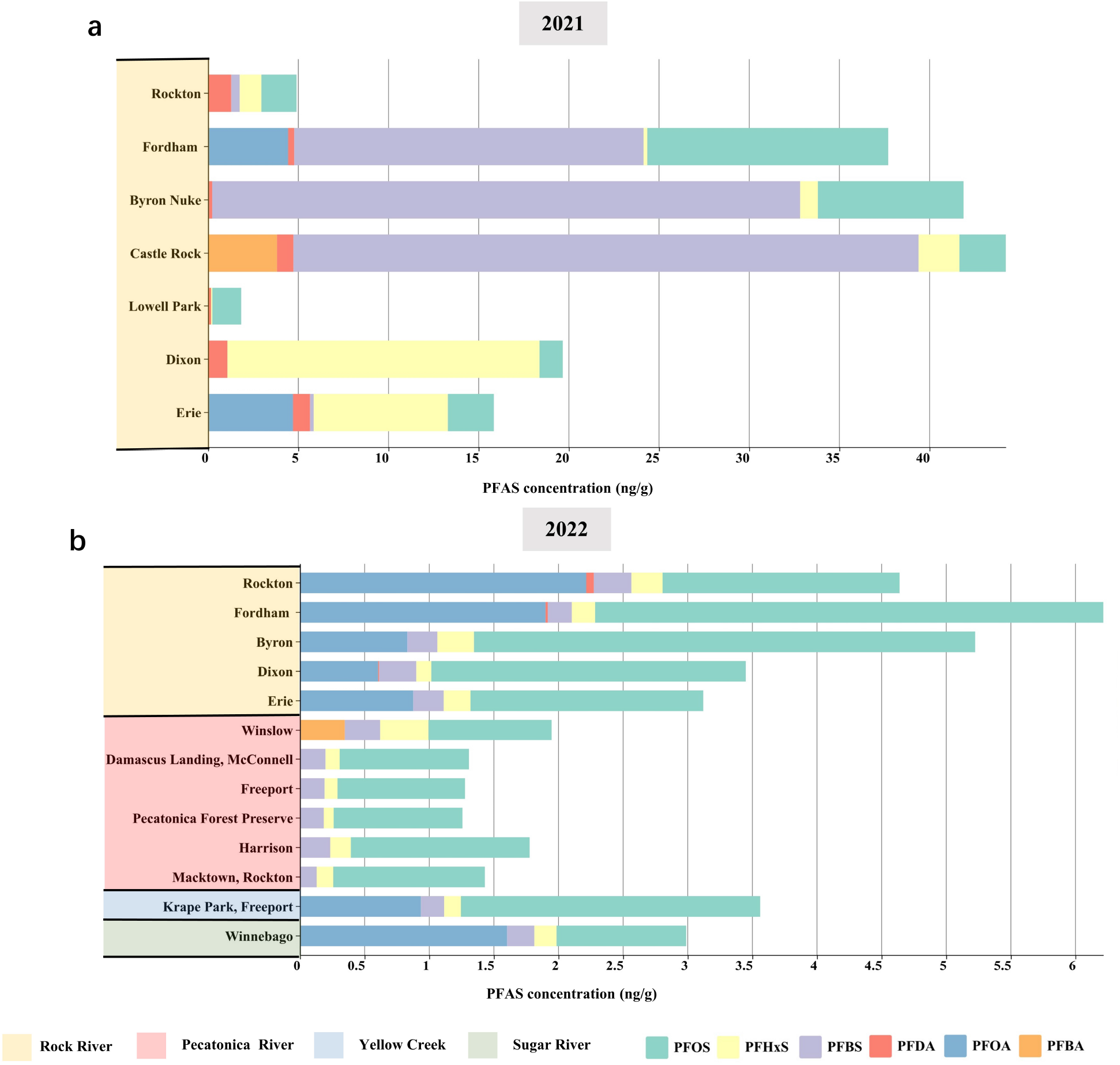
(a). Concentration and composition of PFAS in samples from Rock River (2021). * *Two outlier fish samples, P34-3 and P34-4, from Castle Rock, were excluded from the analysis due to their outlier concentrations of measured PFBA*. **(b)**. Concentration and composition of PFAS in samples from four rivers in Illinois (2022).

Similar PFAS distribution patterns were observed in both years’ fish samples from Rock River sampling sites **(Figure 2)**. Notably, samples from upstream Rockford, IL (Rockton), exhibited lower PFAS concentrations per fish compared to samples from areas closer to urban and industrial activities, such as Rockford, IL, and its immediate downstream areas (Fordham dam, Byron nuclear plant, Castle Rock State Park). Sampling sites further downstream (Lowell Park, Dixon, and Erie) showed decreased PFAS concentrations. This contamination distribution could be influenced by the industries and urban area of Rockford, IL, as it is one of Illinois’ largest cities.

In 2022, among the 17 assessed PFAS compounds, PFHxA demonstrated the highest accumulation (**Figure S2d, e**). The maximum average concentration per fish of 15.42 ng/g was found in samples obtained from Krape Park located in Freeport, followed by PFOS with a maximum concentration per fish of 3.93 ng/g found in samples from the Fordham Dam location on the Rock River located in Rockford. Higher mean concentrations were observed in larger cities like Freeport on the Pecatonica River and Rockford on the Rock River, relative to smaller communities on the same rivers. Higer concentrations are possibly influenced by various factors related to the sources of PFAS contamination and dispersal patterns influenced by human activities, such as urban runoff, especially in areas with high population density or industrial activity (Barbo et al., 2023). Conversely, 8:2 FTS and EtFOSA were not detected in any of the sites, indicating their absence or below detection levels.

These findings are in line with previous studies that also report PFHxA and PFOS as commonly detected PFAS in surface water surveys with PFOS measured at the highest concentrations (Bai and Son, 2021; East et al., 2021; Johnson, 2022). However, the present finds a higher concentration of PFHxA than PFOS in the fish samples, potentially reflecting differing rates of bioaccumulation in wild fish populations, difference due to variations in the sources and pathways of PFAS contamination in the waterways sampled, or the need for multi-timepoint sampling throughout the year. Only bioaccumulation was measured in this study and not direct surface water contamination.

The concentrations of individual PFAS can vary widely even within the same site, possibly due to differences in their chemical properties and their distribution in the environment (Nguyen et al., 2020). For example, at the Winslow site in 2022, the PFAS concentration in the collected samples ranges from non-detectable levels for PFHpA to over 4 ng/g for PFPeA (Table S7). Certain PFAS types are more prevalent in specific sites. PFHxA, PFHxS, PFOS, and PFBS were present in all sites in 2022. Meanwhile, PFBA is only present in samples from Winslow and PFNA is only present in samples from Winnebago. Among the sites, samples from Dixon exhibited the highest diversity, with 12 types of PFAS detected, followed by Rockton, Fordham, and Macktown with 10 types each in 2022. Conversely, Harrison and Damascus Landing had the lowest diversity, with only 5 types of PFAS detected in their samples, likely also a reflection of total numbers of fish sampled (Table 3 and S7). Additionally, the percentage composition of PFAS observed differed across sites in 2022. For instance, in Freeport, PFHxA comprised 77.53% of the total PFAS detected, whereas in Damascus Landing, it accounted for 21.25%. In Forest Preserve, MeFOSA constituted 36.35% of the total PFAS detected, followed by PFHxA and PFOS, contributing 30.07% and 22.73%, respectively, in the fish samples (Table S7).

The differing types of PFAS detected in the same sampling sites between 2021 and 2022 may be the result of a complex interplay between environmental, anthropogenic, and methodological factors. Natural variations in environmental conditions between 2021 and 2022 can impact the mobility, transport, and degradation of PFAS compounds. Different pollution sources, industrial activities, and human behavior can introduce a variety of PFAS compounds into the environment. The presence and types of PFAS compounds can also be influenced by the composition of aquatic ecosystems, including the diversity of species and their behaviors. The overall trend of total PFAS detected from the upstream sampling site at Rockton to the downstream sampling site at Erie remains consistent.

In summary, the concentration and types of PFAS detected in fish samples collected from the 15 sites in Illinois vary. The PFAS with the highest concentrations are PFHxA and PFOS, respectively. The percentage of PFAS in each site also varies, with PFHxA, PFHxS, PFOS and PFBS, and PFOSA being the most common. Sampling locations near higher density communities tend to exhibit higher PFAS contamination. Similarly, larger waterways flowing through more industrialized areas have relatively higher PFAS contamination than smaller waterways.

### 3.2 Comparison of PFAS in fish species

PFAS are persistent in the environment and can accumulate in the food chain, and several studies have noted its accumulation in fish. **Figure 3** and Table S6 and Table S7 presents data on the content and PFAS compounds present in 9 fish types, including Bluegill, Smallmouth Buffalo, Common Carp, Sauger, Smallmouth Bass, Walleye, Black Crappie, Channel Catfish, and Northern Pike, collected from various water bodies in Illinois. The number of individual fish collected and analyzed at each site can be found in **Table 2 and 3**.

**Table 2.** Study sites and sample collection information for the year 2021.

| No. | Location | Waterbody | Date | CCF | SMB | BLG | CAP | WAE | Total (n) |
| --- | --- | --- | --- | --- | --- | --- | --- | --- | --- |
| 1 | Rockton, Winnebago | Rock River | 11.10.2021 | 2 | 2 | 2 | 2 | 2 | 10 |
| 2 | Dixon, Lee | Rock River | 10.15.2021 | 2 | 1 | 1 | 1 | 1 | 6 |
| 3 | Lowell Park, Lee | Rock River | 11.03.2021 | 0 | 2 | 2 | 0 | 0 | 4 |
| 4 | Castle Rock, Ogle | Rock River | 11.05.2021 | 2 | 2 | 2 | 2 | 0 | 8 |
| 5 | Byron Nuclear Plant, Ogle | Rock River | 10.20.2021 | 2 | 3 | 0 | 2 | 1 | 8 |
| 6 | Erie, Whiteside | Rock River | 10.14.2021 | 3 | 1 | 1 | 2 | 1 | 8 |
| 7 | Fordham, Winnebago | Rock River | 11.09.2021 | 1 | 2 | 1 | 2 | 2 | 8 |
| **Total (n)** |  |  |  | 12 | 13 | 9 | 11 | 7 | 52 |

**Table 3.** Study sites and sample collection information for the year 2022.

| No. | Location | Waterbody | Date | BLG | SAB | CAP | SAR | SMB | WAE | BLC | CCF | NOP | Total (n) |
| --- | --- | --- | --- | --- | --- | --- | --- | --- | --- | --- | --- | --- | --- |
| 1 | Rockton, Winnebago | Rock River | 11.03.2022 | 1 | 0 | 2 | 1 | 1 | 0 | 2 | 2 | 1 | 10 |
| 2 | Dixon, Lee | Rock River | 11.07.2022 | 2 | 0 | 1 | 0 | 2 | 2 | 1 | 2 | 0 | 10 |
| 3 | Erie, Whiteside | Rock River | 11.01.2022 | 0 | 0 | 2 | 0 | 2 | 0 | 0 | 3 | 0 | 7 |
| 4 | Fordham, Winnebago | Rock River | 11.10.2022 | 1 | 1 | 2 | 0 | 1 | 2 | 1 | 2 | 0 | 10 |
| 5 | Downstream Byron, Ogle | Rock River | 11.09.2022 | 0 | 0 | 2 | 0 | 2 | 0 | 0 | 1 | 0 | 5 |
| 6 | Harrison, Winnebago | Pecatonica River | 07.28.2022 | 0 | 0 | 2 | 0 | 0 | 0 | 0 | 1 | 0 | 3 |
| 7 | Winslow, Stephenson | Pecatonica River | 08.26.2022 | 0 | 0 | 2 | 0 | 0 | 0 | 0 | 1 | 0 | 3 |
| 8 | Freeport, Stephenson | Pecatonica River | 08.31.2022 | 0 | 0 | 2 | 0 | 0 | 0 | 0 | 1 | 0 | 3 |
| 9 | Macktown, Winnebago | Pecatonica River | 07.28.2022 | 1 | 0 | 2 | 0 | 0 | 2 | 1 | 1 | 0 | 7 |
| 10 | Pecatonica Forest Preserve | Pecatonica River | 08.31.2022 | 0 | 0 | 2 | 0 | 0 | 0 | 0 | 1 | 0 | 3 |
| 11 | Damascus Landing, McConnell | Pecatonica River | 08.26.2022 | 0 | 0 | 2 | 0 | 0 | 0 | 0 | 1 | 0 | 3 |
| 12 | Krape Park, Freeport | Yellow Creek | 08.17.2022 | 0 | 0 | 0 | 0 | 1 | 0 | 0 | 1 | 0 | 2 |
| 13 | Winnebago | Sugar River | 08.01.2022 | 0 | 0 | 2 | 0 | 0 | 0 | 0 | 2 | 0 | 4 |
| **Total (n)** |  |  |  | 5 | 1 | 23 | 1 | 9 | 6 | 5 | 19 | 1 | 70 |

**Figure 3.**
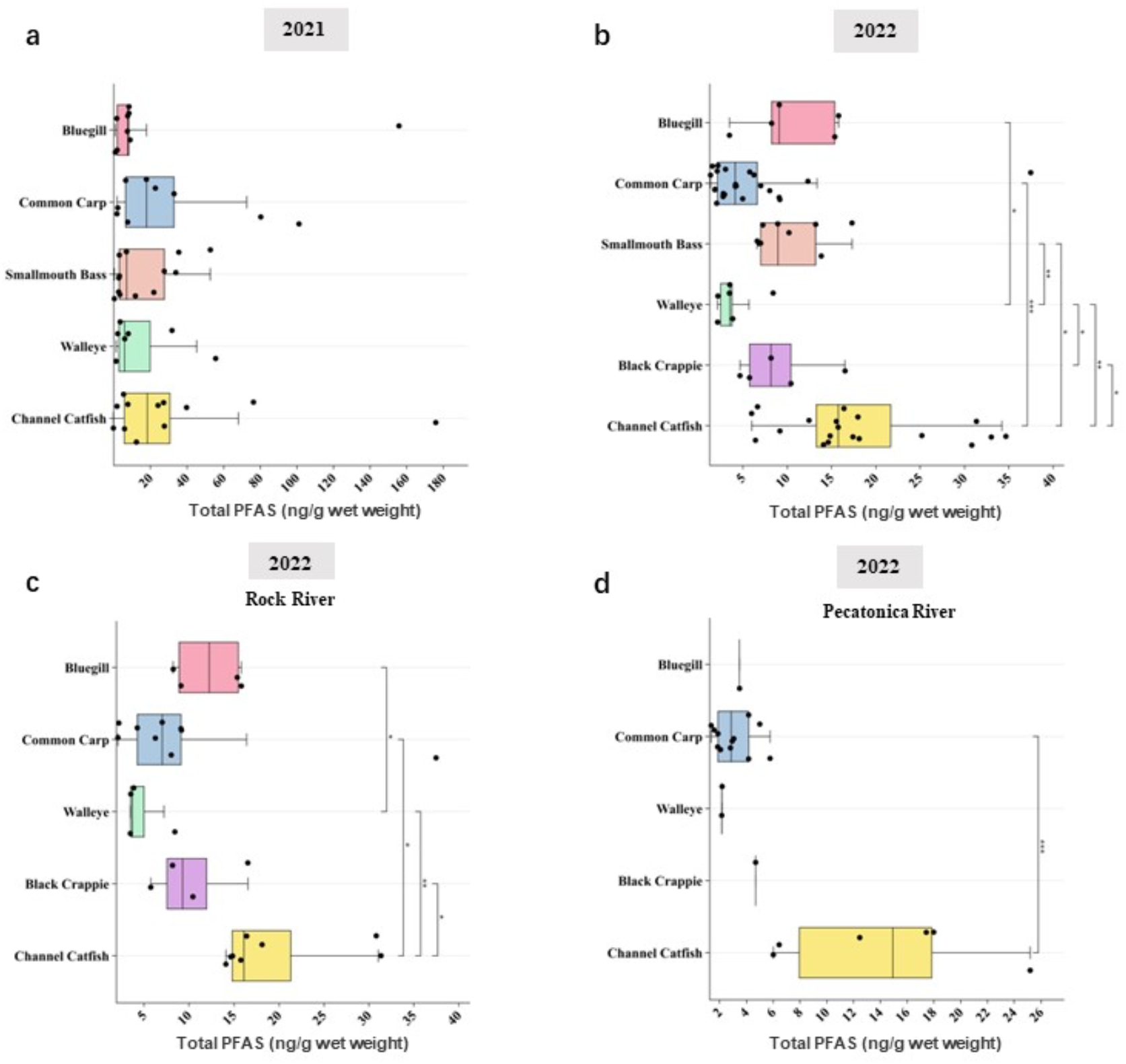
(a). Box-whisker plot of the concentration of total PFAS (PFOS, PFHxS, PFBS, PFHxA, PFOA, PFDA, and PFBA) in five fish species in 2021. *Two outlier fish samples, P34-3 and P34-4, from Castle Rock, were excluded from the analysis due to their outlier concentrations of measured PFBA. (b). Box-whisker plot of the concentration of total PFAS (17 PFAS) in nine fish species in 2022. (c) Box-whisker plot of the concentration of total PFAS (17 PFAS) in five fish species in 2022 in Rock River. (d) Box-whisker plot of the concentration of total PFAS (17 PFAS) in five fish species in 2022 in Pecatonica River.

**Figure 3a and 3b** shows that the Channel Catfish, the highest predator fish included in the study, has the highest total PFAS content in both 2021 and 2022, potentially indicating that PFAS can accumulate in order based on the food chain level. In 2022, fish of various species have consistently higher levels of PFAS accumulation when taken from the Rock River compared to other waterways (Figure 3d, 3e, and Table S10). The greatest differences in waterway source of sample are for the lower trophic species Bluegill and Carp followed by higher trophic species Walley and Crappie. For Channel Catfish, total PFAS contamination is similar but independent of the source waterway (Table S10). Upon comparing data from both 2021 and 2022, it becomes evident that Channel Catfish serves as one of the endpoints for PFAS accumulation. The differences in PFAS contamination patterns between the larger Rock River and the smaller Pecatonica River for lower trophic level fish but not for Channel Catfish could be explained by differences in exposure pathways through diet or direct contact with contaminated water or soils. Further environmental sampling would be required to explore these differences.

In terms of the range of PFAS detected, our study showed that fish in the higher trophic levels have greater number of individual PFAS compounds. Channel Catfish shows a relatively high diversity of the types of PFAS, with 13 types of PFAS detected. This indicates that the fish caught could have been exposed to multiple sources of PFAS exposure. In contrast, Bluegill and Common Carp showed the lowest diversity, with only 11 types of PFAS detected in each fish. With PFHxA accounting for the most significant proportion of all PFAS, we analyzed PFAS other than PFHxA as shown in Figure S3. These results showed no significant cumulative effect of PFAS at different food chain levels after excluding PFHxA data, and their correlation coefficient was found to be very high. Thus, it is likely that PFHxA easily accumulates in the body mass of organisms and is transmitted through the food chain.

### 3.3 Potential risk of PFAS

The Risk Quotient method has been widely used to evaluate the ecological risk of organic pollutants to aquatic ecosystems. The ecotoxicology data of PFAS were obtained from the EPAECTOX database (2022) and the predicted non-effect concentration (PNEC) was calculated. Table S5 summarizes the PNEC data of PFAS. The measured PFAS concentration was divided by the PNEC value to obtain the RQs value of PFAS for fish at each sampling site in Illinois.

The analysis presented in **Figure 4** suggests that PFHxS and PFOS exhibit a notably higher propensity for accumulation in fish across all 13 sampling points within the four waterways in 2022, with a 100% occurrence of high-risk levels. This heightened accumulation potential raises concern on the potential adverse effects on fish health, environmental integrity, and human health. Similarly, PFHxA demonstrates a prevalent risk of accumulation across a majority of sampling sites in this study. Conversely, PFAS compounds such as PFBA, PFHpA, and PFOSA pose low or negligible risks to fish contamination. Notably, PFBS shows high-risk levels across all sampling sites in the Rock River, while being categorized as medium risk in the other three water bodies. This variation could be attributed to differences in the sources of PFBS within different water bodies and influenced by factors such as pH, temperature, and dissolved organic matter (Lyu and Brusseau, 2020; Nguyen et al., 2020). Further research is needed to understand the factors that influence the behavior of PFAS in different water bodies and its transport.

**Figure 4.**
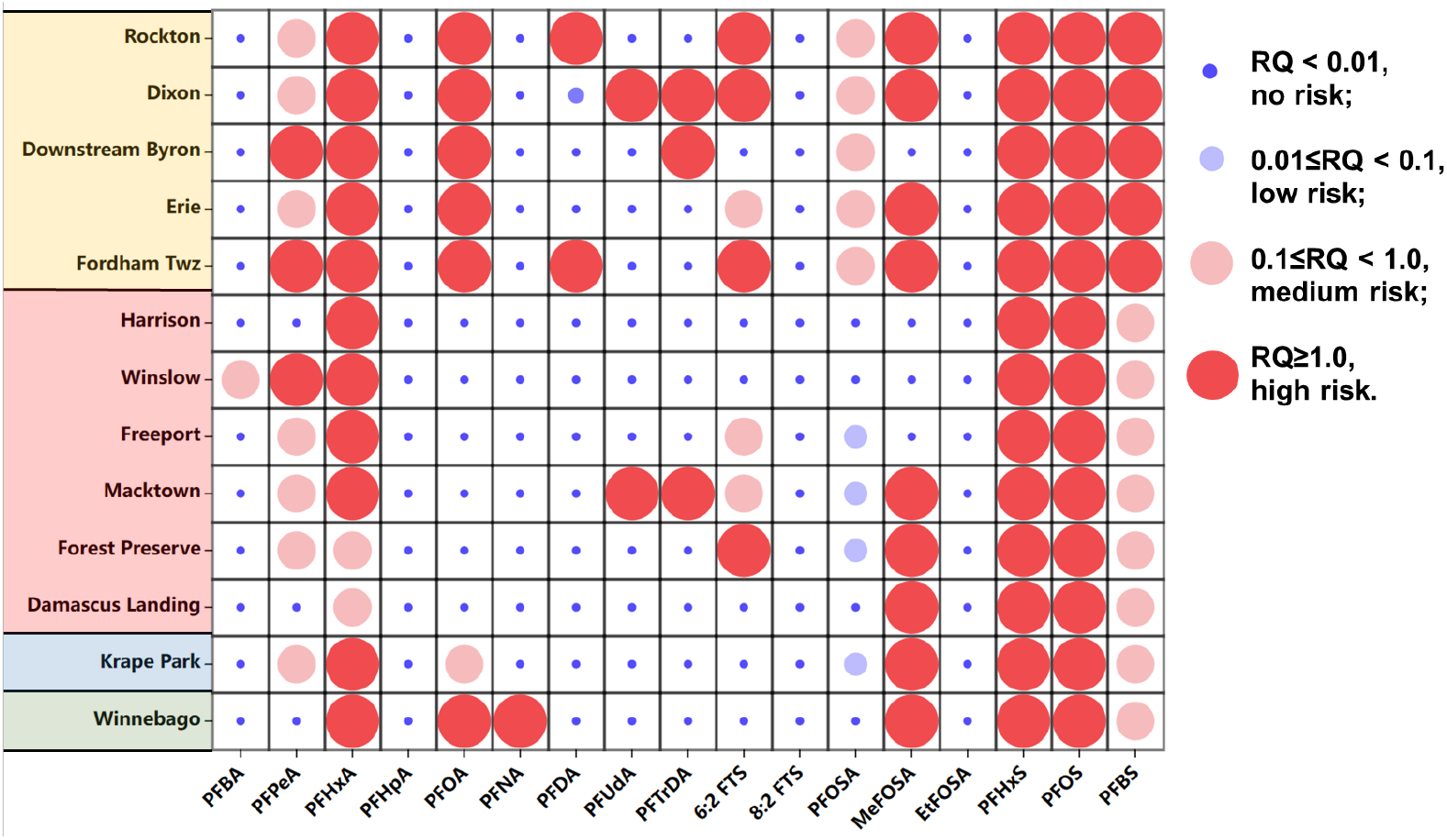
Ecological risk of PFAS from the analysis of fish samples (2022).

Among water bodies, Rock River had the highest number (up to 11) of high-risk PFAS species as evaluated from the samples analyzed. This was followed by Pecatonica River with eight species. The Yellow River had the lowest number of high-risk PFAS, with only four types. In Rock River, up to five PFAS (PFHxA, PFHxS, PFOS, PFOA and PFBS) were identified as high risk at all sampling points, while only two PFAS (PFHxS and PFOS) were identified as high risk at all sampling points in Pecatonica River. The potential risks associated with PFAS exposure are of growing concern, and the findings of this study provide important insights into exposure risks of PFAS in wild caught fish in Illinois. Our results indicate that certain PFAS compounds could pose a significant risk upon consumption, with PFHxS and PFOS being the most toxic.

### 3.4 Fish consumption advisories

In Illinois, the Department of Public Health has established fish consumption advisory triggers for several PFAS compounds, including PFBS, PFHxA, PFOS, PFNA, PFOA, and PFHxS on https://epa.illinois.gov/. These advisory triggers are based on the US Environmental Protection Agency’s RfD and are intended to provide guidance to consumers on safe levels of fish consumption (Cordner et al., 2019).

**Table 4** shows the fish consumption advisory triggers for PFAS in Illinois, based on the frequency of fish consumption and the RfD for each compound. The advisory triggers are given in units of nanograms per gram (ng/g) of fish tissue. According to the advisory triggers, individuals who consume fish daily should limit their consumption of fish containing PFAS. The average levels of PFAS across all nine fish species and 16 sites were above the advisory triggers for safe fish consumption. The level of advisory ranged from consuming fish no more than once a week to no more than once a year. The advisory triggers were mainly based on concentrations of PFOS.

**Table 4.** Illinois PFAS fish Consumption Advisory Triggers (ng/g) (Goodrow et al., 2020a)

| Consumption frequency (meal) | PFBS | PFHxA | PFOS | PFNA | PFOA | PFHxS |
| --- | --- | --- | --- | --- | --- | --- |
| RfD (ng/kg/day) | 320 | 83000 | 1.7 | 3.3 | 2.7 | 16 |
| Unlimited (daily) | ≤98.68 | ≤25594.71 | ≤0.52 | ≤1.02 | ≤0.83 | ≤4.93 |
| Weekly | ≤690.7 | ≤179163 | ≤3.67 | ≤7.12 | ≤5.83 | ≤34.54 |
| Monthly | ≤2960 | ≤767841 | ≤156 | ≤30 | ≤25 | ≤148 |
| Once every 3 months | ≤8881 | ≤2303524 | ≤47 | ≤91.59 | ≤74.93 | ≤444 |
| Yearly | ≤36017 | ≤9342070 | ≤191 | ≤371 | ≤304 | ≤1801 |

| Do not eat | >36017 | >9342070 | >191 | >371 | >304 | >1801 |
| --- | --- | --- | --- | --- | --- | --- |

In conclusion, the fish consumption advisory triggers for PFAS in Illinois provide important guidance to consumers on safe levels of fish consumption. By limiting the consumption of fish containing PFAS to levels below the advisory triggers, consumers can help protect their health and reduce exposure to these harmful chemicals. However, ongoing research and monitoring are needed to better understand the risks of PFAS exposure and to develop more effective strategies for managing these contaminants in the environment.

### 3.5 Policy implication

Based on the data provided, a need for regulation and monitoring of PFAS levels in water sources is imminent. Given the potential risks associated with PFAS exposure, a statewide monitoring program to track PFAS levels in the environment and in fish and wildlife, such as the PFAS Strategic Roadmap released by U.S. EPA in 2021 could be proposed. This could help identify problem areas and inform future policy decisions.

The high levels of PFAS detected in some fish samples suggest that there may be health risks associated with the consumption of certain types of fish. Guidance and advisories for fish consumption are important to inform the public on the potential risks to inform on the safe levels of consumption. It may be beneficial to initiate education and outreach programs to inform the public about the risks associated with PFAS exposure and the potential health effects of consuming PFAS contaminated fish. There is also a need to raise awareness on the safe methods for disposing PFAS-containing products. Overall, the goal should be to minimize the amount of PFAS that enters the environment and to mitigate the health risks associated with exposure to such chemicals. By working together to develop and implement effective policies and strategies, we can protect the health and well-being of both current and future generations.

## 4. Conclusions

Overall, the present study adds to the existing literature on PFAS contamination in fish by providing an inclusive analysis of PFAS levels and composition in the four waterways in Illinois. The findings have important implications for public health and environmental policy, as PFAS contamination in fish can have adverse effects on human health and the ecosystem. The present study highlights the need for continued monitoring and regulation of PFAS contamination in aquatic environments, particularly in areas where the levels of these compounds are high.

The analysis of PFAS contamination in fish samples collected in Illinois during the year 2021 and 2022 reveals significant variations in distribution, concentration, and ecological risks associated with these persistent pollutants. The highest concentrations of PFAS were consistently found in samples from the Rock River, particularly in sites near urban and industrial locations. Notably, PFHxA emerged as the most highly accumulated compound in 2022, while PFBS dominated the concentrations in 2021, suggesting variations in contamination profiles over time and the assess environmental factors that may have contributed to the broad dispersion of PFAS (Langberg et al., 2022). Furthermore, the analysis also revealed distinct trends in PFAS distribution among different fish species and trophic levels, with Channel Catfish consistently exhibiting the highest PFAS content, indicating bioaccumulation potential across the food chain. Of particular concern, PFHxS and PFOS emerged as compounds with notably higher accumulation potentials, raising concerns on adverse effects on fish health and environmental integrity. Given their persistence and potential threat to human health, the elevated levels of PFAS observed across almost all fish species and locations in our study might necessitate fish consumption advisories based on the Illinois PFAS RfD. Our study indicates with certainty the presence of these compounds in fish and the associated health effect based on accumulation. Given the numerous potential sources of PFAS, further studies are warranted to comprehensively evaluate the occurrence and sources of PFAS throughout the state of Illinois. Such information is crucial to better understand the distribution and potential risks of these compounds to the environment.

## Author Contributions

Conceptualization, J.I.; methodology, X.Z., M.S., and M.L; software, X.Z. and M.S.; validation, M.L. and X.Z.; data curation, X.Z., M.S.; writing—original draft preparation, X.Z., and M.S; writing—review and editing, methodology, J.I. and X.Z., M.S, and, T.J.; supervision, J.I.; project administration, J.I. All authors have read and agreed to the published version of the manuscript.

## Institutional Review Board Statement

“Not applicable”

## Informed Consent Statement

“Not applicable.”

## Data Availability Statement

“Not applicable”

## Acknowledgments

Partial funding from IDNR is greatly appreciated. We thank Brenda Koester for valuable insights and discussion.

