## Supplementary Information for "PFAS assessment in fish – samples from Illinois waters"

### SUPPLEMENTARY MATERIALS

### Contents list

Figure S1-S3 and Table S1-S10

**Table S1 Compound,** Abbreviation, and chemical structure of 17 PFASs

| No. | Analyte | Abbreviation | Number of Carbon | Chemical Structure |
|---|---|---|---|---|
| 1 | Perfluoro-butanoic acid | PFBA | C4 | 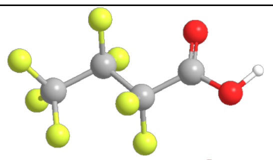 |
| 2 | Perfluoropentanoic acid | PFPeA | C5 | 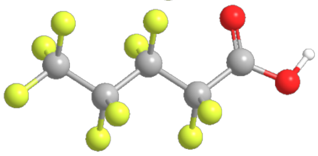 |
| 3 | Perfluoro-hexanoic acid | PFHxA | C6 | 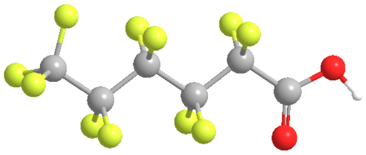 |
| 4 | Perfluoroheptanoic acid | PFHpA | C7 | 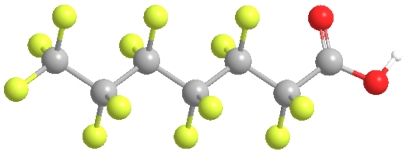 |
| 5 | Perfluoro-octanoic acid | PFOA | C8 | 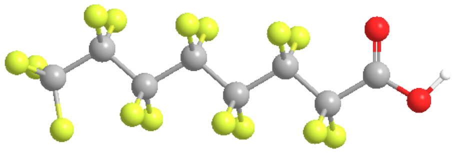 |
| 6 | Perfluorononanoic acid | PFNA | C9 | 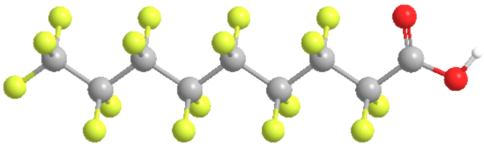 |
| 7 | Perfluoro-decanoic acid | PFDA | C10 | 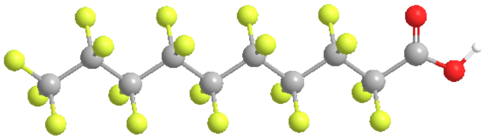 |
| 8 | Perfluoroundecanoic acid | PFUdA | C11 | 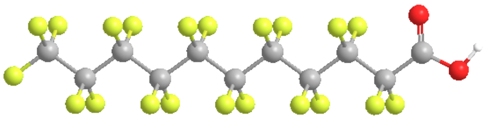 |
| 9 | Perfluorotridecanoic acid | PFTrDA | C13 | 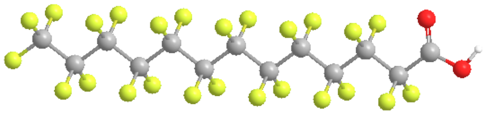 |
| 10 | 6:2 Fluorotelomer sulfonate | 6-2 FTS | C14 | 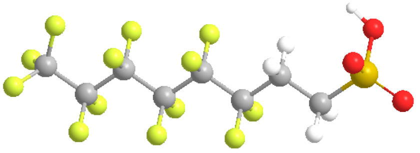 |
| 11 | 8:2 Fluorotelomer sulfonate | 8-2 FTS | C14 | 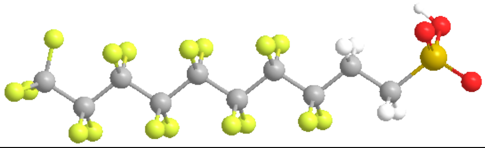 |

| | | | | |
|---|---|---|---|---|
| 12 | Perfluoro-butane sulfonic acid | PFBS | C4 | 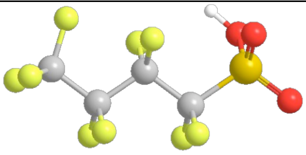 |
| 13 | Perfluoro-hexane sulfonic acid | PFHxS | C6 | 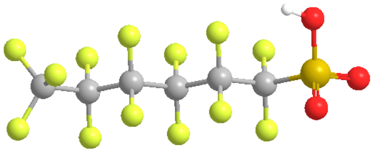 |
| 14 | Perfluoro-octane sulfonic acid | PFOS | C8 | 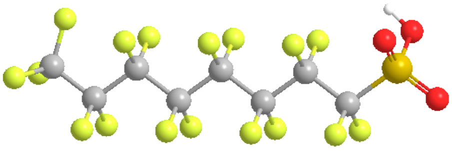 |
| 15 | N-Methyl perfluorooctane sulphonamide | N-MeFOSA | C8 | 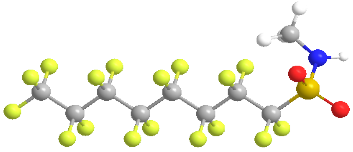 |
| 16 | N-ethylperfluorooctane sulfonamide | N-EtFOSA | C8 | 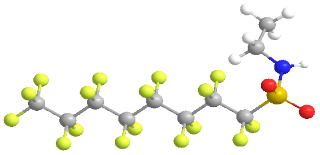 |
| 17 | Perfluorooctane sulfonamide | PFOSA | C8 | 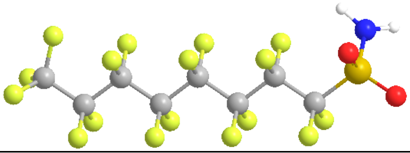 |

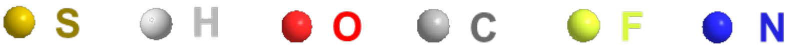

**Table S2** Sample site and site description (2021 and 2022)

| No. | Waterbody | County | Location | Date | Description |
|---|---|---|---|---|---|
| 1 | Rock River | Winnebago | Rockton | 11.03.2022<br>11.10.2021 | This is where Pecatonica River joins Rock River. Rockton is located near the border of Illinois and Wisconsin and has a population of 7,781. |
| 2 | Rock River | Lee | Dixon | 11.07.2022<br>10.15.2021 | Dixon has a population of 15,380 and is home to the Dixon Municipal Airport. The Dixon Fire Department and the Dixon Rural Fire Department are on either side of Rock River. Dixon Iron & Metal Co. significantly contributed to industrial waste but closed in 2019. St Marys Cement-Dixon Plant is located right near Rock River. |
| 3 | Rock River | Lee | Lowell Park | 11.03.2021 | This location is just upstream of Dixon City. |
| 4 | Rock River | Whiteside | Erie | 11.01.2022<br>10.14.2021 | Erie is a village with a population of 1,511. The main industry of Erie is agriculture. |
| 5 | Rock River | Winnebago | Fordham twz | 11.10.2022<br>11.09.2021 | Fordham is a dam located on Rock River. It was built in the Mid 1800s and originally used for hydropower. |
| 6 | Rock River | Ogle | Downstre am Byron | 11.09.2022 | Byron City has a population of 3,775. The main industry is manufacturing, with a focus on precision machining, metal fabrication. |
| 7 | Rock River | Ogle | Byron Leaf River | 10.20.2021 | Leaf River joins Rock River from the west between Byron and Oregon. |
| 8 | Rock River | Ogle | Castle Rock | 11.05.2021 | Castle Rock is a state park. |
| 9 | Pecatonica River | Winnebago | Harrison | 07.28.2022 | With only a population of 934, Harrison is the upstream portion of Pecatonica River before it joins Rock River in Rockton. It is also the location where Sugar River joins Pecatonica River. |
| 10 | Pecatonica River | Stephenson | Winslow | 08.26.2022 | Winslow is a village with a population of 278. It is at the border of Illinois and Wisconsin where Pecatonica River flows southeast into Harrison. |
| 11 | Pecatonica River | Stephenson | Freeport | 08.31.2022 | Freeport has a population of 23,650. It is downstream of Winslow but upstream of Harrison. The city experienced flooding of Pecatonica River the summer of 2022. |
| 12 | Pecatonica River | Winnebago | Macktow n | 07.28.2022 | Macktown Forest Preserve is at the intersection where Pecatonica River meets Rock River. |
| 13 | Pecatonica River | Pecatonica | Forest Preserve | 08.31.2022 | A protected natural area. |

| | | | | | |
|---|---|---|---|---|---|
| 14 | Pecatonica River | Stephenson | Damascus Landing | 08.26.2022 | Damascus is located downstream of Winslow and upstream of Macktown. |
| 15 | Yellow Creek | Stephenson | Krape park | 08.17.2022 | Yellow Creek runs through Krape Park and originates in Freeport from Pecatonica River. There have been many instances of flooding. |
| 16 | Sugar River | Winnebago | Winnebag o | 08.01.2022 | Sugar River is a tributary of Pecatonica River. It joins the Pecatonica River in Winnebago County. |

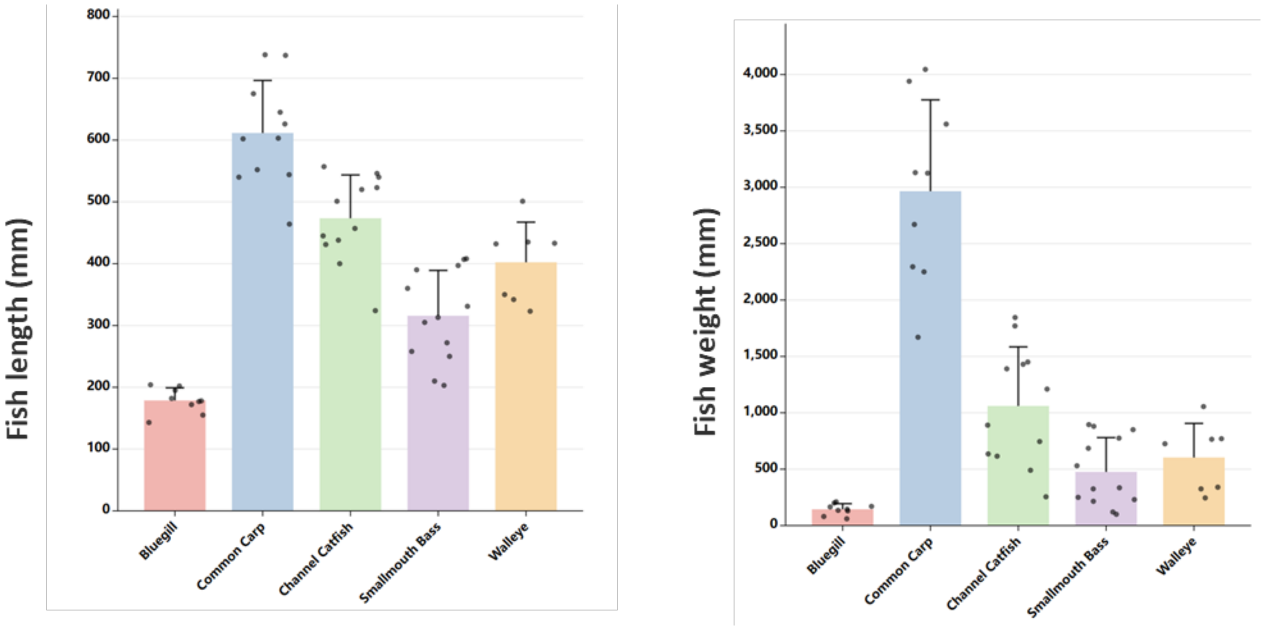


**Figure S1** Length and weight of fish samples collected (2021)

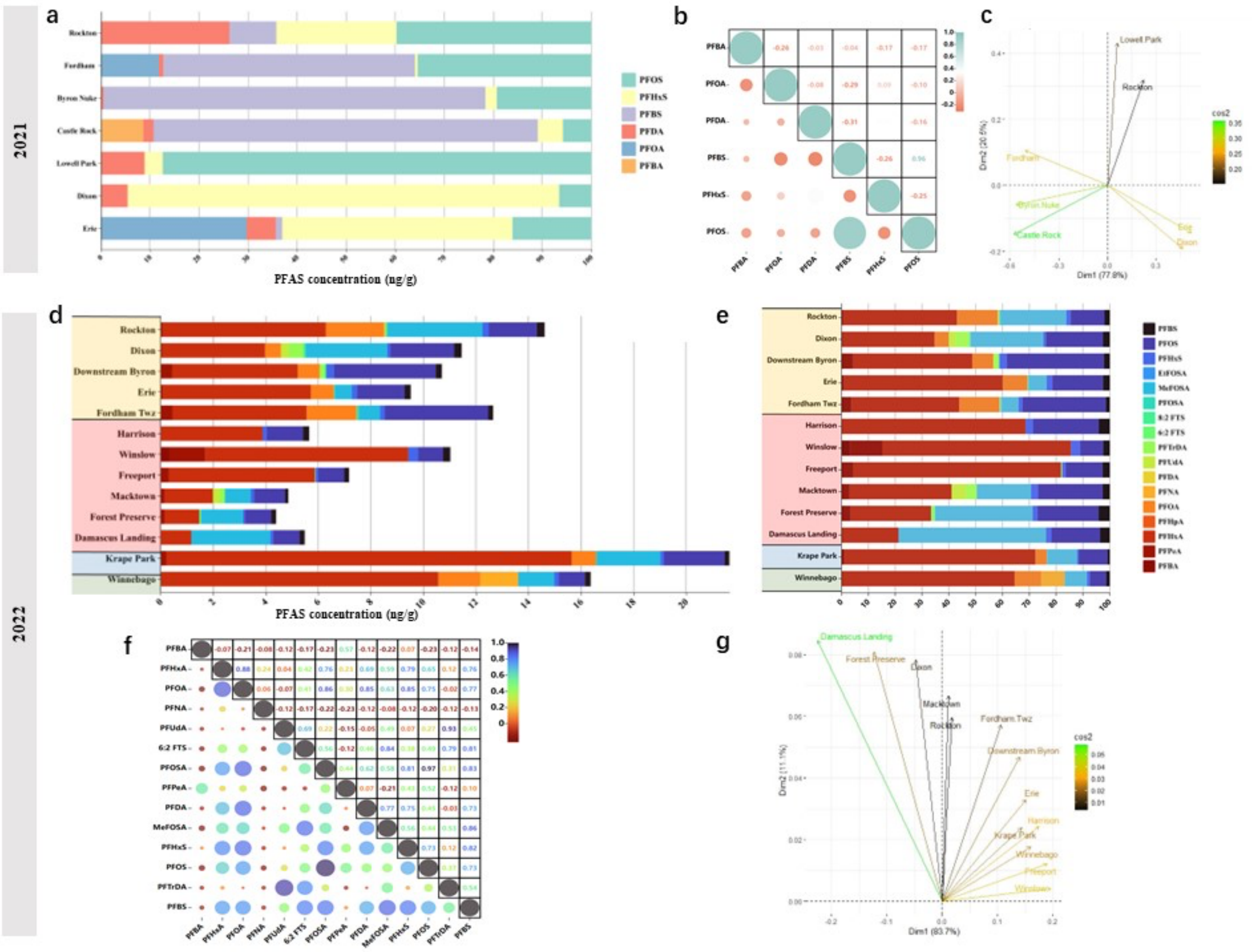


**Figure S2** (a) Mean percentage composition of different PFAS in samples from 7 sites in 2021. ** Two outlier fish samples, P34-3 and P34-4, from Castle Rock, were excluded from the analysis due to their outlier concentrations of measured PFBA.* (b) Correlation analyses between individual PFAS in 2021. (c) Principal Component Analysis (PCA) score across samples obtained from sampling sites in 2021. (d) Concentration and composition of 17 PFAS in samples from Rock River in 2022. (e) Mean percentage composition of different PFAS in samples from 13 sites in 2022. (f) Correlation analyses between individual PFAS in 2022. (c) Principal Component Analysis (PCA) score across samples obtained from sampling sites in 2022.

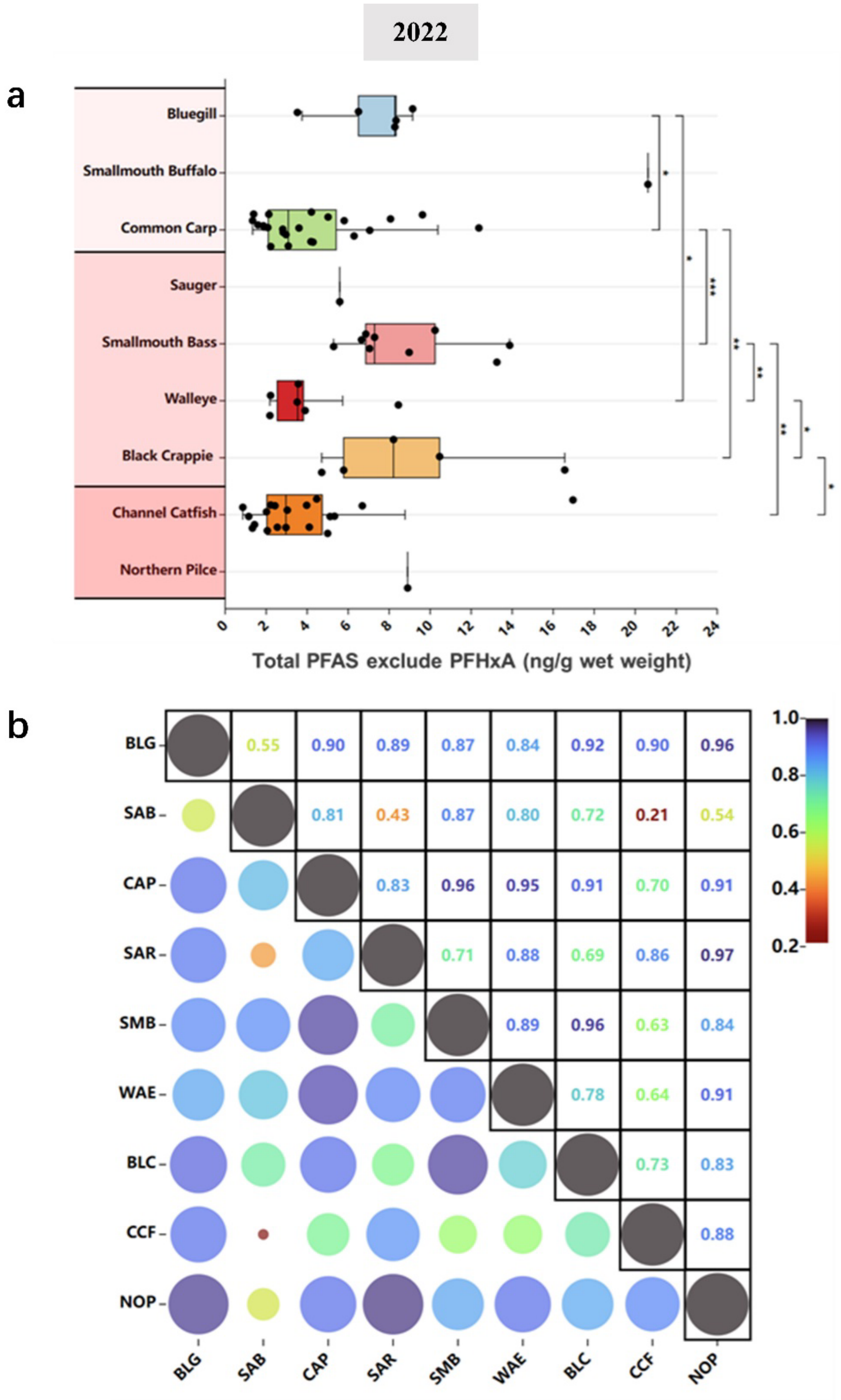


**Figure S3** (a). Box-whisker plot of PFAS (exclude PFHxA) concentration in nine fish species (2022). (b) Correlation matrix of PFAS (exclude PFHxA) for the nine fish species (2022).

**Table S3** Principal Component Analysis (PCA) score for four components across sampling sites (2021)

| Number | Location | Y1 | Y2 | Y3 |
|---|---|---|---|---|
| 1 | Rockton | 0.18 | 0.52 | 0.65 |
| 2 | Fordham | -0.42 | 0.17 | -0.19 |
| 3 | Byron Nuke | -0.47 | -0.09 | 0.22 |
| 4 | Castle Rock | -0.48 | -0.24 | 0.31 |
| 5 | Lowell Park | 0.05 | 0.70 | -0.17 |
| 6 | Dixon | 0.39 | -0.31 | 0.52 |
| 7 | Erie | 0.43 | -0.23 | -0.32 |

**Table S4** Principal Component Analysis (PCA) score for four components across sampling sites (2022)

| Number | Location | Y1 | Y2 | Y3 | Y4 |
|---|---|---|---|---|---|
| 1 | Rockton | 0.03 | 0.31 | 0.26 | 0.59 |
| 2 | Dixon | -0.09 | 0.41 | 0.01 | 0.00 |
| 3 | Downstream.Byron | 0.27 | 0.25 | -0.55 | -0.06 |
| 4 | Erie | 0.29 | 0.17 | 0.04 | 0.13 |
| 5 | Fordham.Twz | 0.20 | 0.30 | -0.47 | 0.39 |
| 6 | Harrison | 0.33 | 0.13 | -0.04 | -0.26 |
| 7 | Winslow | 0.37 | 0.02 | 0.26 | -0.28 |
| 8 | Freeport | 0.36 | 0.06 | 0.19 | -0.23 |
| 9 | Macktown | 0.02 | 0.35 | -0.10 | -0.39 |
| 10 | Forest Preserve | -0.24 | 0.43 | 0.05 | -0.23 |
| 11 | Damascusn Landing | -0.43 | 0.45 | 0.25 | -0.08 |
| 12 | Krape Park | 0.28 | 0.13 | 0.30 | -0.02 |
| 13 | Winnebago | 0.31 | 0.09 | 0.37 | 0.25 |

**Table S5** Predicted Non-Effect Concentration (PNEC) of PFAS

| PNEC(mg/L) | PFBA | PFPeA | PFHxA | PFOA | PFNA | PFDA | PFUdA | PFDoA | PFTrDA | 6-2 FTS |
|---|---|---|---|---|---|---|---|---|---|---|
| Fish | 0.022 | 0.0022 | 0.0088 | 0.0023 | 0.00015 | 0.00015 | 0.00015 | 0.00015 | 0.000013 | 0.000077 |
| PNEC(mg/L) | 8-2 FTS | PFBS | PFHxS | PFOS | N-MeFOSA | N-EtFOSA | PFOSA | PFDS | PFTeDA | |
| Fish | 0.000077 | 0.0011 | 0.000019 | 0.000014 | 0.0011 | 0.0013 | 0.0012 | 0.00016 | 0.0000041 | |

**Table S6** Concentration of 7 PFAS (ng/g) in fish samples at 7 study sites in 2021

| Name | Location | Waterbody | Date | Fish Code | Fish Name | PFOA | PFBS | PFOS | PFDA | PFBA | PFHxS | PFHxA |
|---|---|---|---|---|---|---|---|---|---|---|---|---|
| P05-1 | Rockton | Rock River | 11.10.2021 | CAP | Common Carp | < 0 | < 0 | 0.83 | 1.36 | < 0 | 0.28 | < 0 |
| P05-2 | Rockton | Rock River | 11.10.2021 | CAP | Common Carp | < 0 | < 0 | 4.38 | 1.82 | < 0 | 0.56 | < 0 |
| P05-3 | Rockton | Rock River | 11.10.2021 | CCF | Channel Catfish | < 0 | 4.68 | 1.9 | 0.8 | < 0 | 5.24 | < 0 |
| P05-4 | Rockton | Rock River | 11.10.2021 | CCF | Channel Catfish | < 0 | < 0 | 0.38 | 1.66 | < 0 | < 0 | < 0 |
| P05-5 | Rockton | Rock River | 11.10.2021 | WAE | Walleye | < 0 | < 0 | 3.32 | 2.33 | < 0 | 0.6 | < 0 |
| P05-6 | Rockton | Rock River | 11.10.2021 | SMB | Smallmouth Bass | < 0 | < 0 | 2.54 | 0.46 | < 0 | < 0 | < 0 |
| P05-7 | Rockton | Rock River | 11.10.2021 | SMB | Smallmouth Bass | < 0 | < 0 | 1.3 | 1.38 | < 0 | 0.22 | < 0 |
| P05-8 | Rockton | Rock River | 11.10.2021 | WAE | Walleye | < 0 | < 0 | 1.95 | 0.48 | < 0 | < 0 | < 0 |
| P05-9 | Rockton | Rock River | 11.10.2021 | BLG | Bluegill | < 0 | < 0 | 1.92 | 1.53 | < 0 | 5.1 | < 0 |
| P05-10 | Rockton | Rock River | 11.10.2021 | BLG | Bluegill | < 0 | < 0 | 0.94 | 1 | < 0 | < 0 | < 0 |
| P10-1 | Dixon | Rock River | 10.15.2021 | CAP | Common Carp | < 0 | < 0 | 2.45 | 1.22 | < 0 | 14.4 | < 0 |
| P10-2 | Dixon | Rock River | 10.15.2021 | SMB | Smallmouth Bass | < 0 | < 0 | 1.62 | 1.6 | < 0 | 3.95 | < 0 |
| P10-3 | Dixon | Rock River | 10.15.2021 | BLG | Bluegill | < 0 | < 0 | 2.16 | 1.11 | < 0 | 5.91 | < 0 |
| P10-4 | Dixon | Rock River | 10.15.2021 | CCF | Channel Catfish-Large | < 0 | < 0 | 0.59 | 0.89 | < 0 | 75 | < 0 |
| P10-5 | Dixon | Rock River | 10.15.2021 | CCF | Channel Catfish-Small | < 0 | < 0 | 0.06 | 1.16 | < 0 | 4.37 | < 0 |
| P10-6 | Dixon | Rock River | 10.15.2021 | WAE | Walleye | < 0 | < 0 | 0.88 | 0.42 | < 0 | 0.19 | < 0 |
| P20-1 | Lowell Park | Rock River | 11.03.2021 | SMB | Smallmouth Bass-Lg | < 0 | < 0 | 0.58 | < 0 | < 0 | < 0 | < 0 |
| P20-2 | Lowell Park | Rock River | 11.03.2021 | SMB | Smallmouth Bass-Sm | < 0 | < 0 | 3.08 | 0.19 | < 0 | 0.27 | < 0 |

| | | | | | | | | | | | | |
|---|---|---|---|---|---|---|---|---|---|---|---|---|
| P20-3 | Lowell Park | Rock River | 11.03.2021 | BLG | Bluegill-Sm | < 0 | < 0 | 1.06 | < 0 | < 0 | < 0 | < 0 |
| P20-4 | Lowell Park | Rock River | 11.03.2021 | BLG | Bluegill-Lg | < 0 | < 0 | 1.68 | 0.46 | < 0 | < 0 | < 0 |
| P34-1 | Castle Rock | Rock River | 11.05.2021 | SMB | Smallmouth Bass-Sm | < 0 | 10 | 5.89 | 0.58 | 22.9 | 13.6 | < 0 |
| P34-2 | Castle Rock | Rock River | 11.05.2021 | SMB | Smallmouth Bass-Lg | < 0 | 6.08 | 5.05 | 0.92 | < 0 | < 0 | < 0 |
| P34-3 | Castle Rock | Rock River | 11.05.2021 | CAP | Common Carp-Lg | < 0 | 7.18 | 1.07 | 1.26 | 175 | 0.25 | 6.45 |
| P34-4 | Castle Rock | Rock River | 11.05.2021 | CAP | Common Carp-Sm | < 0 | 138 | 0.89 | 1.05 | 285 | 0.36 | 8.63 |
| P34-5 | Castle Rock | Rock River | 11.05.2021 | CCF | Channel Catfish-Sm | < 0 | 173 | 1.99 | 0.99 | < 0 | < 0 | < 0 |
| P34-6 | Castle Rock | Rock River | 11.05.2021 | CCF | Channel Catfish-Lg | < 0 | 6.98 | 0.11 | 0.86 | < 0 | < 0 | < 0 |
| P34-7 | Castle Rock | Rock River | 11.05.2021 | BLG | Bluegill-Lg | < 0 | 5.3 | 1.31 | 1.24 | < 0 | < 0 | < 0 |
| P34-8 | Castle Rock | Rock River | 11.05.2021 | BLG | Bluegill-Sm | < 0 | 6.6 | 1.06 | 0.89 | < 0 | < 0 | < 0 |
| P40-1 | Byron Nuke Discharge Leaf River Delta | Rock River | 10.20.2021 | SMB | Smallmouth Bass-Sm | < 0 | 32.4 | 1.75 | < 0 | < 0 | < 0 | < 0 |
| P40-2 | Byron Nuke Discharge Leaf River Delta | Rock River | 10.20.2021 | SMB | Smallmouth Bass-Md | < 0 | 32.2 | 2.92 | 0.32 | < 0 | 0.28 | < 0 |
| P40-3 | Byron Nuke Discharge Leaf River Delta | Rock River | 10.20.2021 | SMB | Smallmouth Bass-Lg | < 0 | 25.7 | 1.64 | < 0 | < 0 | 0.48 | < 0 |
| P40-4 | Byron Nuke Discharge Leaf River Delta | Rock River | 10.20.2021 | CCF | Channel Catfish-Sm | 0.125 | 36.8 | 1.4 | < 0 | < 0 | 1.67 | < 0 |
| P40-5 | Byron Nuke Discharge Leaf River Delta | Rock River | 10.20.2021 | CCF | Channel Catfish-Lg | < 0 | 27.6 | 0.26 | < 0 | < 0 | < 0 | < 0 |
| P40-6 | Byron Nuke Discharge Leaf River Delta | Rock River | 10.20.2021 | WAE | Walleye | < 0 | 39.4 | 14.1 | < 0 | < 0 | 2.35 | < 0 |

| | | | | | | | | | | | | |
|---|---|---|---|---|---|---|---|---|---|---|---|---|
| P40-7 | Byron Nuke Discharge Leaf River Delta | Rock River | 10.20.2021 | CAP | Common Carp-Sm | < 0 | 24.7 | 6.34 | 1.16 | < 0 | 0.97 | < 0 |
| P40-8 | Byron Nuke Discharge Leaf River Delta | Rock River | 10.20.2021 | CAP | Common Carp-Lg | 0.18 | 42 | 36.2 | < 0 | < 0 | 2.1 | < 0 |
| P46-1 | Erie | Rock River | 10.14.2021 | SMB | Smallmouth Bass | 14.1 | < 0 | 7.37 | 0.66 | < 0 | < 0 | < 0 |
| P46-2 | Erie | Rock River | 10.14.2021 | BLG | Bluegill | 3.21 | 0.72 | 2.46 | 1.29 | < 0 | < 0 | < 0 |
| P46-3 | Erie | Rock River | 10.14.2021 | WAE | Walleye | 1.17 | < 0 | 5.53 | 0.2 | < 0 | 1.25 | < 0 |
| P46-4 | Erie | Rock River | 10.14.2021 | CCF | Channel Catfish | 17.5 | < 0 | 1.95 | 1.63 | < 0 | 3.33 | < 0 |
| P46-5 | Erie | Rock River | 10.14.2021 | CCF | Channel Catfish | 1.65 | 0.93 | 0.17 | 0.93 | < 0 | 2.42 | < 0 |
| P46-6 | Erie | Rock River | 10.14.2021 | CCF | Channel Catfish | < 0 | < 0 | 0.65 | 1.01 | < 0 | 25.8 | < 0 |
| P46-7 | Erie | Rock River | 10.14.2021 | CAP | Common Carp | < 0 | < 0 | 1.15 | 1.2 | < 0 | 20.6 | < 0 |
| P46-8 | Erie | Rock River | 10.14.2021 | CAP | Common Carp | < 0 | < 0 | 1.12 | 0.64 | < 0 | 6.09 | < 0 |
| P87-1 | Fordham | Rock River | 11.09.2021 | BLG | Bluegill | < 0 | 155 | 0.9 | < 0 | < 0 | < 0 | < 0 |
| P87-2 | Fordham | Rock River | 11.09.2021 | CAP | Common Carp-Lg | 6.09 | < 0 | 91.2 | 2.67 | < 0 | 1.38 | < 0 |
| P87-3 | Fordham | Rock River | 11.09.2021 | CAP | Common Carp-Sm | < 0 | < 0 | 2.02 | < 0 | < 0 | < 0 | < 0 |
| P87-4 | Fordham | Rock River | 11.09.2021 | CCF | Channel Catfish | < 0 | < 0 | < 0 | 0.04 | < 0 | < 0 | < 0 |
| P87-5 | Fordham | Rock River | 11.09.2021 | SMB | Smallmouth Bass | < 0 | < 0 | 3.34 | < 0 | < 0 | < 0 | < 0 |
| P87-6 | Fordham | Rock River | 11.09.2021 | WAE | Walleye | < 0 | < 0 | 3.74 | < 0 | < 0 | < 0 | < 0 |
| P87-7 | Fordham | Rock River | 11.09.2021 | WAE | Walleye | 28.4 | < 0 | 3.63 | < 0 | < 0 | < 0 | < 0 |
| P87-8 | Fordham | Rock River | 11.09.2021 | SMB | Smallmouth Bass | 0.985 | < 0 | 2.04 | < 0 | < 0 | 0.22 | < 0 |

**Table S7** Concentration of 17 PFAS (ng/g) in fish samples at 13 study sites in 2022

| Name | Location | Waterbody | Date | Fish Code | Fish Name | PFBA | PFPeA | PFHxA | PFHpA | PFOA | PFNA | PFDA | PFUdA | PFTrDA | 6:2 FTS | 8:2 FTS | PFOSA | MeFOSA | EtFOSA | PFHxS | PFOS | PFBS |
|---|---|---|---|---|---|---|---|---|---|---|---|---|---|---|---|---|---|---|---|---|---|---|
| P05-1 | Rockton | Rock River | 11.03.2022 | CCF | Channel Catfish | <0 | 0.63 | <0 | <0 | <0 | <0 | <0 | <0 | <0 | <0 | <0 | 0.03 | 3.98 | <0 | 0.54 | 1.07 | 0.43 |
| P05-2 | Rockton | Rock River | 11.03.2022 | CCF | Channel Catfish | <0 | <0 | 5.11 | <0 | <0 | <0 | 0.59 | <0 | <0 | <0 | <0 | 0.01 | 2.81 | <0 | 0.18 | 0.35 | 0.17 |
| P05-3 | Rockton | Rock River | 11.03.2022 | SAR | Sauger(Hg) | <0 | <0 | 23.74 | <0 | <0 | <0 | <0 | <0 | <0 | <0 | <0 | 0.07 | 3.18 | <0 | 0.09 | 1.94 | 0.31 |
| P05-4 | Rockton | Rock River | 11.03.2022 | | Northern Pilce | <0 | <0 | <0 | <0 | 1.02 | <0 | <0 | <0 | <0 | 0.06 | <0 | 0.27 | 4.20 | <0 | 0.06 | 3.17 | 0.12 |
| P05-5 | Rockton | Rock River | 11.03.2022 | BLC | Black Crappie (Hg) | <0 | <0 | <0 | <0 | 1.81 | <0 | <0 | <0 | <0 | <0 | <0 | 0.03 | 1.19 | <0 | 0.24 | 2.25 | 0.26 |
| P05-6 | Rockton | Rock River | 11.03.2022 | BLG | Bluegill | <0 | <0 | <0 | <0 | 2.95 | <0 | <0 | <0 | <0 | <0 | <0 | 0.01 | 4.02 | <0 | 0.47 | 1.33 | 0.36 |
| P05-7 | Rockton | Rock River | 11.03.2022 | SMB | Smallmouth Bass | <0 | <0 | <0 | <0 | 5.31 | <0 | <0 | <0 | <0 | 0.08 | <0 | 0.06 | 3.89 | <0 | 0.25 | 3.89 | 0.38 |
| P05-8 | Rockton | Rock River | 11.03.2022 | CAP | Common Carp | <0 | <0 | 5.53 | <0 | <0 | <0 | <0 | <0 | <0 | 0.07 | <0 | <0 | 1.81 | <0 | 0.25 | 1.16 | 0.32 |
| P05-9 | Rockton | Rock River | 11.03.2022 | CAP | Common Carp | <0 | <0 | 27.88 | <0 | 3.06 | <0 | <0 | <0 | <0 | 0.06 | <0 | <0 | 5.03 | <0 | 0.14 | 1.11 | 0.22 |
| P05-10 | Rockton | Rock River | 11.03.2022 | BLC | Black Crappie (Hg) | <0 | <0 | <0 | <0 | 7.99 | <0 | <0 | <0 | <0 | 0.09 | <0 | 0.02 | 5.87 | <0 | 0.19 | 2.07 | 0.34 |
| P10-1 | Dixon | Rock River | 11.07.2022 | CAP | Common Carp | <0 | <0 | <0 | <0 | <0 | <0 | <0 | <0 | <0 | 0.11 | <0 | <0 | 1.76 | <0 | 0.08 | 2.01 | 0.32 |
| P10-2 | Dixon | Rock River | 11.07.2022 | WAE | Walleye | <0 | <0 | <0 | <0 | <0 | <0 | <0 | <0 | <0 | 0.08 | <0 | 0.14 | 3.23 | <0 | 0.06 | 4.68 | 0.27 |
| P10-3 | Dixon | Rock River | 11.07.2022 | WAE | Walleye | <0 | <0 | <0 | <0 | <0 | <0 | <0 | <0 | <0 | <0 | <0 | 0.13 | 2.01 | <0 | 0.08 | 1.33 | 0.34 |
| P10-4 | Dixon | Rock River | 11.07.2022 | CCF | Channel Catfish | <0 | <0 | 9.01 | <0 | <0 | <0 | <0 | <0 | <0 | 0.12 | <0 | <0 | 3.98 | <0 | <0 | 0.61 | 0.41 |
| P10-5 | Dixon | Rock River | 11.07.2022 | CCF | Channel Catfish | <0 | <0 | 13.90 | <0 | <0 | <0 | <0 | 2.93 | 5.32 | 0.08 | <0 | 0.02 | 6.95 | <0 | 0.24 | 1.02 | 0.39 |
| P10-6 | Dixon | Rock River | 11.07.2022 | BLC | Black Crappie (Hg) | <0 | <0 | <0 | <0 | <0 | <0 | <0 | <0 | <0 | 0.14 | <0 | <0 | 4.21 | <0 | 0.09 | 3.31 | 0.45 |

| | | | | | | | | | | | | | | | | | | | | | | |
|---|---|---|---|---|---|---|---|---|---|---|---|---|---|---|---|---|---|---|---|---|---|---|
| P10-7 | Dixon | Rock River | 11.07.2022 | BLG | Bluegill | <0 | 0.40 | 7.07 | <0 | 2.27 | <0 | <0 | <0 | <0 | 0.05 | <0 | <0 | 3.87 | <0 | 0.21 | 1.27 | 0.26 |
| P10-8 | Dixon | Rock River | 11.07.2022 | BLG | Bluegill | <0 | <0 | 9.34 | <0 | 1.77 | <0 | <0 | <0 | <0 | 0.06 | <0 | 0.02 | 1.79 | <0 | 0.16 | 2.60 | 0.11 |
| P10-9 | Dixon | Rock River | 11.07.2022 | SMB | Smallmouth Bass | <0 | <0 | <0 | <0 | <0 | <0 | 0.07 | <0 | <0 | 0.04 | <0 | 0.09 | 1.78 | <0 | 0.09 | 4.48 | 0.10 |
| P10-10 | Dixon | Rock River | 11.07.2022 | SMB | Smallmouth Bass | <0 | <0 | <0 | <0 | 2.01 | <0 | <0 | <0 | <0 | 0.04 | <0 | 0.04 | 1.38 | <0 | 0.16 | 3.02 | 0.22 |
| P40-1 | Downstream Byron | Rock River | 11.09.2022 | CCF | Channel Catfish | <0 | 0.98 | 11.81 | <0 | 1.87 | <0 | <0 | <0 | <0 | <0 | <0 | <0 | <0 | <0 | 0.27 | 0.55 | 0.31 |
| P40-2 | Downstream Byron | Rock River | 11.09.2022 | SMB | Smallmouth Bass | <0 | <0 | 12.05 | <0 | <0 | <0 | <0 | <0 | <0 | <0 | <0 | 0.08 | <0 | <0 | 0.21 | 4.82 | 0.18 |
| P40-3 | Downstream Byron | Rock River | 11.09.2022 | SMB | Smallmouth Bass | <0 | 0.68 | <0 | <0 | <0 | <0 | <0 | <0 | 1.04 | <0 | <0 | 0.13 | <0 | <0 | 0.34 | 4.72 | 0.14 |
| P40-4 | Downstream Byron | Rock River | 11.09.2022 | CAP | Common Carp | <0 | 0.58 | <0 | <0 | <0 | <0 | <0 | <0 | <0 | <0 | <0 | 0.04 | <0 | <0 | 0.19 | 5.28 | 0.20 |
| P40-5 | Downstream Byron | Rock River | 11.09.2022 | CAP | Common Carp | <0 | <0 | <0 | <0 | 2.28 | <0 | <0 | <0 | <0 | <0 | <0 | 0.01 | <0 | <0 | 0.41 | 4.02 | 0.33 |
| P46-1 | Erie | Rock River | 11.01.2022 | CAP | Common Carp Large | <0 | <0 | <0 | <0 | <0 | <0 | <0 | <0 | <0 | 0.07 | <0 | 0.00 | <0 | <0 | 0.15 | 1.65 | 0.26 |
| P46-2 | Erie | Rock River | 11.01.2022 | CCF | Channel Catfish-Medium | <0 | <0 | 12.13 | <0 | 1.39 | <0 | <0 | <0 | <0 | <0 | <0 | 0.00 | <0 | <0 | 0.16 | 0.78 | 0.22 |
| P46-3 | Erie | Rock River | 11.01.2022 | CAP | Common Carp Small | <0 | <0 | <0 | <0 | <0 | <0 | <0 | <0 | <0 | <0 | <0 | 0.01 | <0 | <0 | 0.21 | 1.86 | 0.15 |
| P46-4 | Erie | Rock River | 11.01.2022 | CCF | Channel Catfish-Large | <0 | <0 | 14.01 | <0 | <0 | <0 | <0 | <0 | <0 | <0 | <0 | 0.01 | 1.89 | <0 | 0.11 | 0.27 | 0.15 |
| P46-5 | Erie | Rock River | 11.01.2022 | SMB | Smallmouth Bass Small | <0 | <0 | <0 | <0 | <0 | <0 | <0 | <0 | <0 | <0 | <0 | 0.21 | 2.53 | <0 | 0.19 | 5.86 | 0.18 |
| P46-6 | Erie | Rock River | 11.01.2022 | CCF | Channel Catfish-Small | <0 | 0.45 | 13.44 | <0 | <0 | <0 | <0 | <0 | <0 | <0 | <0 | <0 | <0 | <0 | 0.46 | 0.22 | 0.31 |
| P46-7 | Erie | Rock River | 11.01.2022 | SMB | Smallmouth Bass-Large | <0 | <0 | <0 | <0 | 4.74 | <0 | <0 | <0 | <0 | <0 | <0 | 0.04 | <0 | <0 | 0.17 | 1.97 | 0.38 |
| P87-1 | Fordham | Rock River | 11.10.2022 | BLG | Bluegill | <0 | 0.89 | <0 | <0 | <0 | <0 | <0 | <0 | <0 | 0.05 | <0 | 0.04 | 2.55 | <0 | 0.34 | 4.21 | 0.20 |
| P87-2 | Fordham | Rock River | 11.10.2022 | SMB | Smallmouth Bass | <0 | 0.43 | <0 | <0 | 3.65 | <0 | <0 | <0 | <0 | <0 | <0 | 0.28 | 4.38 | <0 | 0.17 | 4.17 | 0.18 |

| | | | | | | | | | | | | | | | | | | | | | | |
|---|---|---|---|---|---|---|---|---|---|---|---|---|---|---|---|---|---|---|---|---|---|---|
| P87-3 | Fordham | Rock River | 11.10.2022 | SAB | Smallmouth Buffalo | <0 | 0.93 | <0 | <0 | 2.61 | <0 | 0.17 | <0 | <0 | <0 | <0 | 0.16 | <0 | <0 | 0.26 | 16.29 | 0.21 |
| P87-4 | Fordham | Rock River | 11.10.2022 | BLC | Black Crappie (Hg) | <0 | 0.36 | <0 | <0 | 3.84 | <0 | <0 | <0 | <0 | 0.04 | <0 | 0.06 | <0 | <0 | 0.38 | 5.54 | 0.25 |
| P87-5 | Fordham | Rock River | 11.10.2022 | CCF | Channel Catfish | <0 | 0.48 | 26.05 | <0 | 3.78 | <0 | <0 | <0 | <0 | <0 | <0 | 0.03 | <0 | <0 | 0.30 | 0.50 | 0.24 |
| P87-6 | Fordham | Rock River | 11.10.2022 | CCF | Channel Catfish | <0 | 0.36 | 17.01 | <0 | <0 | <0 | <0 | <0 | <0 | <0 | <0 | 0.03 | <0 | <0 | 0.15 | 0.44 | 0.17 |
| P87-7 | Fordham | Rock River | 11.10.2022 | CAP | Common Carp | <0 | 1.13 | <0 | <0 | 5.12 | <0 | <0 | <0 | <0 | 0.04 | <0 | 0.00 | <0 | <0 | 0.11 | 1.43 | 0.23 |
| P87-8 | Fordham | Rock River | 11.10.2022 | CAP | Common Carp | <0 | <0 | 7.86 | <0 | <0 | <0 | <0 | <0 | <0 | <0 | <0 | 0.00 | <0 | <0 | 0.05 | 1.14 | 0.15 |
| P87-9 | Fordham | Rock River | 11.10.2022 | WAE | Walleye | <0 | <0 | <0 | <0 | <0 | <0 | <0 | <0 | <0 | <0 | <0 | 0.06 | 0.61 | <0 | 0.01 | 2.77 | 0.11 |
| P87-10 | Fordham | Rock River | 11.10.2022 | WAE | Walleye | <0 | <0 | <0 | <0 | <0 | <0 | <0 | <0 | <0 | <0 | <0 | 0.07 | 0.46 | <0 | 0.03 | 2.83 | 0.12 |
| PW01-1 | Harrison | Pecatonica River | 07.28.2022 | CAP | Common Carp | <0 | <0 | <0 | <0 | <0 | <0 | <0 | <0 | <0 | <0 | <0 | <0 | <0 | <0 | 0.15 | 0.91 | 0.33 |
| PW01-2 | Harrison | Pecatonica River | 07.28.2022 | CAP | Common Carp | <0 | <0 | <0 | <0 | <0 | <0 | <0 | <0 | <0 | <0 | <0 | 0.00 | <0 | <0 | 0.19 | 2.68 | 0.21 |
| PW01-3 | Harrison | Pecatonica River | 07.28.2022 | CCF | Channel Catfish | <0 | <0 | 11.63 | <0 | <0 | <0 | <0 | <0 | <0 | <0 | <0 | <0 | <0 | <0 | 0.14 | 0.56 | 0.16 |
| PW02-1 | Winslow | Pecatonica River | 08.26.2022 | CCF | Channel Catfish | <0 | 0.82 | 23.21 | <0 | <0 | <0 | <0 | <0 | <0 | <0 | <0 | <0 | <0 | <0 | 0.56 | 0.34 | 0.29 |
| PW02-2 | Winslow | Pecatonica River | 08.26.2022 | CAP | Common Carp | <0 | 0.87 | <0 | <0 | <0 | <0 | <0 | <0 | <0 | <0 | <0 | 0.00 | <0 | <0 | 0.27 | 1.45 | 0.24 |
| PW02-3 | Winslow | Pecatonica River | 08.26.2022 | CAP | Common Carp | 1.04 | 2.33 | <0 | <0 | <0 | <0 | <0 | <0 | <0 | <0 | <0 | <0 | <0 | <0 | 0.29 | 1.07 | 0.29 |
| PW08-1 | Freeport | Pecatonica River | 08.31.2022 | CCF | Channel Catfish | <0 | 0.93 | 16.68 | <0 | <0 | <0 | <0 | <0 | <0 | <0 | <0 | 0.00 | <0 | <0 | 0.06 | 0.18 | 0.16 |
| PW08-2 | Freeport | Pecatonica River | 08.31.2022 | CAP | Common Carp | <0 | <0 | <0 | <0 | <0 | <0 | <0 | <0 | <0 | 0.03 | <0 | 0.01 | <0 | <0 | 0.13 | 1.23 | 0.21 |
| PW08-3 | Freeport | Pecatonica River | 08.31.2022 | CAP | Common Carp | <0 | <0 | <0 | <0 | <0 | <0 | <0 | <0 | <0 | 0.04 | <0 | 0.00 | <0 | <0 | 0.11 | 1.55 | 0.20 |
| PW11-1 | Macktown | Pecatonica River | 07.28.2022 | CAP | Common Carp | <0 | <0 | <0 | <0 | <0 | <0 | <0 | <0 | <0 | <0 | <0 | 0.00 | 0.83 | <0 | 0.04 | 0.89 | 0.11 |
| PW11-2 | Macktown | Pecatonica River | 07.28.2022 | CAP | Common Carp | <0 | <0 | <0 | <0 | <0 | <0 | <0 | <0 | <0 | 0.03 | <0 | 0.01 | 0.37 | <0 | 0.07 | 1.46 | 0.15 |

| | | | | | | | | | | | | | | | | | | | | | | |
|---|---|---|---|---|---|---|---|---|---|---|---|---|---|---|---|---|---|---|---|---|---|---|
| PW11-3 | Macktown | Pecatonica River | 07.28.2022 | CCF | Channel Catfish | <0 | <0 | 12.99 | <0 | <0 | <0 | <0 | <0 | 1.34 | <0 | <0 | 0.01 | 2.55 | <0 | 0.10 | 0.29 | 0.18 |
| PW11-4 | Macktown | Pecatonica River | 07.28.2022 | WAE | Walleye | <0 | <0 | <0 | <0 | <0 | <0 | <0 | <0 | <0 | <0 | <0 | 0.02 | 1.47 | <0 | 0.05 | 0.56 | 0.08 |
| PW11-5 | Macktown | Pecatonica River | 07.28.2022 | WAE | Walleye | <0 | <0 | <0 | <0 | <0 | <0 | <0 | <0 | <0 | <0 | <0 | 0.02 | 0.86 | <0 | 0.11 | 1.12 | 0.10 |
| PW11-6 | Macktown | Pecatonica River | 07.28.2022 | BLG | Bluegill | <0 | <0 | <0 | <0 | <0 | <0 | <0 | 1.78 | <0 | <0 | <0 | 0.00 | 0.76 | <0 | 0.06 | 0.83 | 0.08 |
| PW11-7 | Macktown | Pecatonica River | 07.28.2022 | BLC | Black Crappie (Hg) | <0 | 0.97 | <0 | <0 | <0 | <0 | <0 | <0 | <0 | <0 | <0 | 0.01 | <0 | <0 | 0.47 | 3.06 | 0.20 |
| PW17-1 | Forest Preserve | Pecatonica River | 08.31.2022 | CCF | Channel Catfish | <0 | <0 | 3.97 | <0 | <0 | <0 | <0 | <0 | <0 | 0.09 | <0 | 0.00 | 1.59 | <0 | 0.13 | 0.06 | 0.19 |
| PW17-2 | Forest Preserve | Pecatonica River | 08.31.2022 | CAP | Common Carp | <0 | <0 | <0 | <0 | <0 | <0 | <0 | <0 | <0 | 0.05 | <0 | 0.00 | 2.49 | <0 | 0.06 | 1.43 | 0.16 |
| PW17-3 | Forest Preserve | Pecatonica River | 08.31.2022 | CAP | Common Carp | <0 | 0.44 | <0 | <0 | <0 | <0 | <0 | <0 | <0 | 0.05 | <0 | 0.01 | 0.72 | <0 | 0.04 | 1.50 | 0.20 |
| PW19-1 | Damascus Landing | Pecatonica River | 08.26.2022 | CCF | Channel Catfish | <0 | <0 | 3.50 | <0 | <0 | <0 | <0 | <0 | <0 | <0 | <0 | <0 | 2.42 | <0 | 0.08 | 0.29 | 0.18 |
| PW19-2 | Damascus Landing | Pecatonica River | 08.26.2022 | CAP | Common Carp | <0 | <0 | <0 | <0 | <0 | <0 | <0 | <0 | <0 | <0 | <0 | <0 | 3.08 | <0 | 0.22 | 2.29 | 0.20 |
| PW19-3 | Damascus Landing | Pecatonica River | 08.26.2022 | CAP | Common Carp | <0 | <0 | <0 | <0 | <0 | <0 | <0 | <0 | <0 | <0 | <0 | <0 | 3.53 | <0 | 0.03 | 0.42 | 0.21 |
| PWN03-1 | Krape Park | Yellow Creek | 08.17.2022 | SMB | Smallmouth Bass | <0 | <0 | <0 | <0 | 1.87 | <0 | <0 | <0 | <0 | <0 | <0 | 0.05 | 3.72 | <0 | 0.06 | 4.35 | 0.18 |
| PWN03-2 | Krape Park | Yellow Creek | 08.17.2022 | CCF | Channel Catfish | <0 | 0.44 | 30.84 | <0 | <0 | <0 | <0 | <0 | <0 | <0 | <0 | 0.01 | 1.11 | <0 | 0.20 | 0.28 | 0.18 |
| PWB03-1 | Winnebago | Sugar River | 08.01.2022 | CCF | Channel Catfish | <0 | <0 | 12.55 | <0 | 1.91 | <0 | <0 | <0 | <0 | <0 | <0 | 0.01 | <0 | <0 | 0.18 | 0.73 | 0.20 |
| PWB03-2 | Winnebago | Sugar River | 08.01.2022 | CCF | Channel Catfish | <0 | <0 | 29.71 | <0 | 2.76 | <0 | <0 | <0 | <0 | <0 | <0 | <0 | <0 | <0 | 0.35 | 1.67 | 0.22 |
| PWB03-3 | Winnebago | Sugar River | 08.01.2022 | CAP | Common Carp | <0 | <0 | <0 | <0 | <0 | <0 | <0 | <0 | <0 | <0 | <0 | <0 | 2.04 | <0 | 0.02 | 0.52 | 0.22 |
| PWB03-4 | Winnebago | Sugar River | 08.01.2022 | CAP | Common Carp | <0 | <0 | <0 | <0 | 1.74 | 5.77 | <0 | <0 | <0 | <0 | <0 | <0 | 3.43 | <0 | 0.14 | 1.09 | 0.20 |

**Table S8.** Identifiers and LC-MS parameters for Target PFAS.

| Name | Abbreviation | CAS Number | Retention Time (min) | Transition | Quantification Reference Compound | LOQ (ng/mL) |
|---|---|---|---|---|---|---|
| Perfluoro-n-butanoic acid | PFBA | 375-22-4 | 11.9 | 213.0>169.0 | 13C4-PFBA | 1.0 |
| Perfluoro-n-pentanoic acid | PFPeA | 2706-90-3 | 12.51 | 262.9>219.0 | 13C5-PFPeA | 0.5 |
| Perfluoro-n-hexanoic acid | PFHxA | 307-24-4 | 12.87 | 313.0>269.0 | 13C5-PFPeA | 0.5 |
| Perfluoro-n-heptanoic acid | PFHpA | 375-85-9 | 13.15 | 363.0>319.0 | 13C4-PFHpA | 1.0 |
| Perfluoro-n-octanoic acid | PFOA | 335-67-1 | 13.46 | 413.0>369.0 | 13C4-PFBA | 1.0 |
| Perfluoro-n-nonanoic acid | PFNA | 375-95-1 | 13.77 | 463.0>419.0 | 13C4-PFBA | 1.0 |
| Perfluoro-n-decanoic acid | PFDA | 335-76-2 | 14.07 | 513.0>469.0 | 13C4-PFBA | 0.05 |
| Perfluoro-n-undecanoic acid | PFUdA | 2058-94-8 | 14.39 | 563.0>519.0 | 13C2-PFDoA | 1.0 |
| Perfluoro-n-dodecanoic acid | PFDoA | 307-55-1 | 14.66 | 613.0>569.0 | 13C2-PFDoA | 1.0 |
| Perfluoro-n-tridecanoic acid | PFTrDA | 72629-94-8 | 14.92 | 663.0>619.0 | 13C2-PFDoA | 0.1 |
| Perfluoro-n-tetradecanoic acid | PFTeDA | 376-06-7 | 15.15 | 713.0>669.0 | 13C2-PFTeDA | 1.0 |
| Perfluorohexadecanoic acid | PFHxDA | 67905-19-5 | 15.14 | 813.0>769.0 | 13C2-PFTeDA | 1.0 |
| Perfluoro-1-butanesulfonate | PFBS | 29420-49-3 | 12.5 | 298.9>80.0 | 13C4-PFOS | 0.05 |
| Perfluorohexane sulfonate | PFHxS | 355-46-4 | 13.1 | 399.0>80.0 | 13C4-PFOS | 0.05 |
| Perfluoro-1-heptane sulfonate | PFHpS | 21934-50-9 | 13.4 | 449.0>80.0 | 13C4-PFOS | 0.05 |
| Perfluoro-n-octanoic sulfonate | PFOS | 1763-23-1 | 13.68 | 499.0>80.0 | 13C4-PFOS | 0.1 |
| Perfluoro-1-decanesulfonate | PFDS | 2806-15-7 | 14.31 | 599.0>80.0 | 13C4-PFOS | 0.05 |
| 1h 1h 2h 2h-perfluorooctanesulfonate | 6:2 FTS | 27619-97-2 | 13.46 | 427.0>407.0 | M2-6:2 FTS | 0.05 |
| 1h 1h 2h 2h- | 8:2 FTS | 27619-96-1 | 14.08 | 527.0>507.0 | M2-8:2 FTS | 0.05 |

| | | | | | | |
|---|---|---|---|---|---|---|
| perfluorodecanesulfonate | | | | | | |
| Perfluoro-1-octanesulfonamide | PFOSA | 754-91-6 | 14.71 | 498.0>78.0 | 13C8-PFOSA | 0.1 |
| n-methylperfluoro-1-octanesulfonamideacetic acid | MeFOSA | 2355-31-9 | 14.28 | 512.0>169.0 | d3MeFOSA | 0.1 |
| n-ethylperfluoro-1-octanesulfonamideacetic acid | EtFOSA | 2991-50-6 | 14.46 | 526.0>169.0 | d5EtFOSA | 0.1 |
| Bis(trifluoromethane)sulfonimide lithium salt | HQ115 | 90076-65-6 | 12.11 | 280.0>146.0 | 13C4-PFOS | 1.0 |

**Table S9.** Identifiers and LC-MS parameters for labeled PFAS internal standards.

| Name | Abbreviation | CAS Number | Retention Time (min) | Transition |
|---|---|---|---|---|
| 13C5-Perfluoro-n-pentanoic acid | 13C5-PFPeA | 2483735-37-9 | 12.51 | 268.0>223.0 |
| 13C4-Perfluoro-n-heptanoic acid | 13C4-PFHpA | 2328024-55-9 | 13.13 | 367.0>322.0 |
| 13C2-Perfluoro-n-dodecanoic acid | 13C2-PFDoA | 960315-52-0 | 14.65 | 615.0>570.0 |
| 13C2-Perfluoro-n-tetradecanoic acid | 13C2-PFTeDA | 2708218-82-8 | 15.13 | 715.0>670.0 |
| 13C4-Perfluoro-n-butanoic acid | 13C4-PFBA | 1017281-29-6 | 11.89 | 217.0>172.0 |
| 13C4-Perfluoro-n-octanoic sulfonate | 13C4-PFOS | 960315-48-4 | 13.7 | 503.0>80.0 |
| M2-1h 1h 2h 2h-perfluorooctanesulfonate | M2-6:2 FTS | 2708218-89-5 | 13.43 | 429.0>409.0 |
| M2-1h 1h 2h 2h-perfluorodecanesulfonate | M2-8:2 FTS | 2708218-90-8 | 14.09 | 529.0>509.0 |
| 13C8-Perfluoro-1-octanesulfonamide | 13C8-PFOSA | 1365803-60-6 | 14.7 | 506.0>78.0 |
| d3-n-methylperfluoro-1-octanesulfonamideacetic acid | d3-MeFOSA | 1400690-70-1 | 14.28 | 515.0>169.0 |

| | | | | |
|---|---|---|---|---|
| d5-n-ethylperfluoro-1-octanesulfonamideacetic acid | d5-EtFOSA | 1265205-97-7 | 14.43 | 531.0>169.0 |

**Table S10.** Comparison of total PFAS concentration (ng/g ww) in five fish species in Rock River versus Non-Rock River in 2022.

| Fish Species | Total | Rock River | Non-Rock River |
|---|---|---|---|
| Bluegill | 10.44 | 12.17 | 3.51 |
| Common Carp | 6.00 | 9.54 | 3.72 |
| Walleye | 3.97 | 4.86 | 2.20 |
| Black Crappie | 9.15 | 10.26 | 4.71 |
| Catfish | 17.96 | 17.22 | 18.78 |

* *Comparisons only made for species with more than 1 sample per waterway.*
